# High-throughput siRNA screening reveals functional interactions and redundancies of human processive exoribonucleases

**DOI:** 10.1101/2020.08.05.238006

**Authors:** Anna Hojka-Osinska, Aleksander Chlebowski, Ewelina P. Owczarek, Kamila Afek, Kamila Kłosowska, Roman J. Szczesny, Andrzej Dziembowski

## Abstract

Processive exoribonucleases, the executors of RNA decay, participate in multiple physical and functional interactions. Unlike physical ones, functional relationships have not been investigated in human cells. Here we have screened cells deficient in DIS3, XRN2, EXOSC10, DIS3L, and DIS3L2 with a custom siRNA library and determined their functional interactions with diverse pathways of RNA metabolism. We uncover a complex network of positive interactions that buffer alterations in RNA degradation. We reveal important reciprocal actions between RNA decay and transcription and explore alleviating interactions between RNA splicing and DIS3 mediated degradation. We also use a large scale library of genes associated with RNA metabolism to determine genetic interactions of nuclear DIS3 and cytoplasmic DIS3L, revealing their unique functions in RNA degradation and uncovering cooperation between the cytoplasmic degradation and nuclear processing of RNA. Finally, genome-wide siRNA screening of DIS3 reveals processes such as microtubule organization and regulation of telomerase activity that are also functionally associated with nuclear exosome-mediated RNA degradation.

## INTRODUCTION

In eukaryotes, gene expression is a very complex process regulated at numerous levels from chromatin structure throughout transcription, pre-RNA processing, RNA localization, up to translation. In the process of transcription, each cell synthesizes an excessive amount of RNA from different classes, and most of them are subjected to extensive processing or degradation. For example, pre-messenger RNAs (pre-mRNAs) that are transcribed by RNA polymerase II (RNAPII) need to undergo maturation process, that involve many steps (*i.e.* addition of cap structure at 5’ end, splicing, 3’end processing) that are tightly interconnected and predominantly cotranscriptional (Moore & Proudfoot, 2009). In addition, most RNAPII promoters are bidirectional, and aside from producing transcripts in the sense orientation (pre-mRNAs), they produce short-lived promoter upstream transcripts (PROMPTs) in the antisense orientation, that are quickly removed (Andersson *et al*, 2015). Furthermore, other independent transcription units produce precursors of several other functional RNA classes (e.g., pre-ribosomal RNA [rRNA], pre-small nuclear RNA [snRNA], and primary micro RNA [pri-miRNA]) and pervasive transcripts, such as long intergenic non-coding RNAs (lincRNA) (Cech & Steitz, 2014). All of them first need to undergo a maturation process that involves a series of modification steps. A plethora of different factors are involved in regulating the post-transcriptional life of RNA molecules. For example, pre-mRNA splicing requires five different snRNAs and more than 100 proteins(Wahl *et al*, 2009). There are more than 60 different RNA helicases that are encoded in the human genome that control mRNA and rRNA biogenesis and decay. Nucleotidyltransferases, such as poly(A) and poly(U) polymerases, add non-templated nucleotides to the 3’ end of RNA molecules and are involved in transfer RNA (tRNA), mRNA, and U6 snRNA maturation and decay. Additionally, stringent quality-control pathways exist at every step, thereby ensuring that aberrant transcripts are quickly degraded (Schmid & Jensen, 2010; Kilchert *et al*, 2016).

Processive exoribonucleases play a major role in eukaryotic RNA turnover and processing. RNA molecules can be degraded or processed from the 5’ end by enzymes that belong to the XRN family of RNases(Nagarajan *et al*, 2013). This is believed to be the case for most RNAs in *Saccharomyces cerevisiae*. In human cells, the 3’-to-5’ direction, which involves the exosome complex or monomeric DIS3-like exonuclease 2 (DIS3L2), appears to be more prevalent (Chlebowski *et al*, 2013, 2010; Lubas *et al*, 2013).The exosome is a large macromolecular assembly that is composed of the nine-subunit, ring-shaped, catalytically inactive core and differentially localized catalytic subunits that belong to the DIS3 and RRP6 protein families (DIS3, DIS3L, and EXOSC10/RRP6 in humans) (Chlebowski *et al*, 2013; Dziembowski *et al*, 2007; Tomecki *et al*, 2010; Lebreton *et al*, 2008). The RNA substrates of exosome complexes that contain DIS3 or DIS3L pass through the central channel of the exosome ring to reach the exoribonuclease active site (Drążkowska *et al*, 2013; Makino *et al*, 2013, 2015; Malet *et al*, 2010). Yeast Rrp6 is positioned at the top of the exosome ring (Wasmuth *et al*, 2014).

The human DIS3 protein is endowed with both exonucleolytic and endonucleolytic activities that reside in the ribonuclease binding (RNB) and PilT N-terminus (PIN) domains, respectively (Tomecki *et al*, 2010; Lebreton *et al*, 2008; Schaeffer *et al*, 2009; Schneider *et al*, 2009). DIS3 is mainly nuclear but excluded from the nucleolus. It is involved in the 3’-end processing of stable RNA species (5.8S rRNA, small nucleolar RNA [snoRNA]) and degrades pervasive transcription products, such as PROMPTs and enhancer RNAs (Chlebowski *et al*, 2013; Tomecki *et al*, 2010, 2014). It is also crucial for nuclear RNA quality control (Kilchert *et al*, 2016; Szczepińska *et al*, 2015). DIS3 inactivation leads to the global dysregulation of gene expression and is lethal as a consequence (Szczepińska *et al*, 2015; Tomecki *et al*, 2014). *DIS3* is one of the most frequently mutated genes in multiple myeloma (MM), which is a bone marrow malignancy that is characterized by diffuse marrow infiltration, focal bone lesions, and soft-tissue (extramedullary) disease. It is fatal and incurable (Prideaux *et al*, 2014; Kumar *et al*, 2017). Mutations of *DIS3* are found in 10-20% of MM patients. This mutation frequency is considerably high, given that MM is not particularly prone to somatic mutations (Lawrence *et al*, 2013). Most mutations are located within the RNB domain and impair the protein’s exonucleolytic activity to some extent but usually do not destroy it completely (Tomecki *et al*, 2014; Chapman *et al*, 2011).

DIS3L has the same domain architecture as DIS3 but exhibits only exoribonucleolytic activity because of mutations in the PIN domain active site. The protein localizes to the cytoplasm where it participates in mRNA turnover in association with the exosome core complex (Lubas *et al*, 2013; Tomecki *et al*, 2010; Staals *et al*, 2010). Information about DIS3L function remains very fragmentary.

Although the exosome complex degrades both single-stranded and partially double-stranded RNA substrates *in vitro*, it needs cofactors to achieve optimal activity and specificity *in vivo*. Various complexes can activate the exosome in the cytoplasm and nucleus, but they all share the common feature of the presence of RNA helicases of the MTR4/SKI2 family.

DIS3L2 is a cytoplasmic 3’-5’ exoribonuclease that to date has not been shown to be a part of any stable macromolecular assembly (Lubas *et al*, 2013). Mutations of *DIS3L2* that affect catalytic activity of the protein are associated with Perlman syndrome, a rare genetic overgrowth disease (Astuti *et al*, 2012). DIS3L2 is known to be involved in the decay of selected mRNAs and pre-miRNAs from the let-7 family (Lubas *et al*, 2013; Chang *et al*, 2013; Ustianenko *et al*, 2013). The DIS3L2-mediated decay of pre-let7 mRNA depends on the addition of non-templated uridines to the 3’ end of the pre-mRNA and plays a crucial role in the maintenance of stem-cell pluripotency (Chang *et al*, 2013; Ustianenko *et al*, 2013). DIS3L2 contributes to the surveillance/decay of medium-sized RNA species(Łabno *et al*, 2016).

EXOSC10 (also known as RRP6) is a poorly processive/distributive exoribonuclease, that works in association with exosome complex. EXOSC10 is predominantly nuclear protein that is enriched in nucleoli and involved in 18S rRNA 3’-end processing (Sloan *et al*, 2013). It also controls the levels of mature snoRNAs (Szczepińska *et al*, 2015; Macias *et al*, 2015) and might assist DIS3 in some nucleoplasmic functions. Furthermore, connections between EXOSC10 and DNA repair have been reported (Marin-Vicente *et al*, 2015).

XRN1 and XRN2are processive 5’-to-3’ exoribonucleases that degrade 5’-monophosphorylated RNA species (Nagarajan *et al*, 2013). XRN1 is a cytoplasmic protein that is involved in bulk mRNA decay, a process that requires mRNA to be decapped by the DCP2/DCP1 complex. XRN2 is localized in the nucleus, but its function is less well described. The best-documented role of XRN2 is its involvement in the termination of RNAPII transcription termination (Eaton *et al*, 2018; Fong *et al*, 2015). Moreover, XRN2 participates in the 5’-end processing of snoRNAs and rRNAs and is implicated in nuclear RNA surveillance (Davidson *et al*, 2012).

In simple and genetically tractable organisms, analyses of relationships between different pathways and the identification of novel factors and pathways are usually conducted through genetic screening. The development of high-throughput methods, such as synthetic genetic array (SGA)(Tong *et al*, 2001)or diploid-based synthetic lethality analysis on microarray (dSLAM)(Pan *et al*, 2007) for mapping and analyzing genetic interactions (GIs), was an important breakthrough for expanding our knowledge of GI networks on a genome-wide scale and broaden research possibilities in functional genomics. The yeast *Saccharomyces cerevisiae* are used for large scale synthetic lethal screens thanks to its advantages in experimental easiness. However, recent advances in gene silencing technology provided an experimental format for high-throughput GI mapping in higher organisms. This approach has already been used in a large number of studies of many basic cellular processes, such as genome maintenance and the regulation of cell division(Mukherji *et al*, 2006; Sokolova *et al*, 2016; Stojic *et al*, 2020; Paulsen *et al*, 2009; Kavanaugh *et al*, 2015), but RNA metabolism has rarely been studied using this kind of approach, with the exception of nonsense-mediated decay (NMD), stress granule/P-body assembly, and ribosomal biogenesis (Ohn *et al*, 2008; Badertscher *et al*, 2015). To our knowledge, no studies have focused on such a fundamental process like RNA decay.

In the present study, we performed multiple high-throughput small interfering RNA (siRNA) screens using cells that were deficient in DIS3, XRN2, EXOSC10, DIS3L, and DIS3L2. All of the models were interrogated with an siRNA collection that targeted 280 genes that are involved in core RNA metabolism processes. We systematically analyzed genes that interact with query ribonucleases and comprise a network that gathers known and novel functional associations, revealing the deep interplay between RNA decay and other RNA metabolism pathways. The observed unique GI profiles of the analyzed ribonucleases indicated that they play unique roles in RNA degradation. Shared GIs suggested some cooperation in the context of such processes as RNA splicing, nuclear export, and RNA transcription. We also performed high-throughput siRNA screens to characterize functional interactions between main cytoplasmic and nuclear exosome-mediated RNA degradation, extending the siRNA library to target 3904 genes that are involved in RNA metabolism. These analyses revealed that DIS3 and DIS3L have divergent functions not only because of different cellular localization and substrate specificity but also at the level of functional associations. Nevertheless, we also observed some redundancy that corroborated RNA nuclear export. Notably, we found a deep interplay between cytoplasmic RNA degradation and nuclear RNA metabolism. Finally, we performed genome-wide siRNA screen covering 18104 human genes. With this approach we derived a network of all possible DIS3 interactions with cellular processes. Our results provide a unique overview of functional associations of DIS3 and offer a discovery tool that can be used to dissect the roles of DIS3 in biological processes beyond RNA decay.

Overall, the present study identified functional interactions of main mammalian ribonucleases with different cellular processes, thereby providing an expansive view of the interplay and coordinated action of different processes that buffer imbalances that are introduced by the dysfunction of RNA degradation.

## RESULTS

### Nuclear exosome cofactors were synthetically lethal with exosome dysfunction, whereas cytoplasmic cofactors had the opposite effect

We sought to identify novel functional interactions between proteins, pathways, and complexes that are involved in RNA metabolism based on high-throughput siRNA screens that involved the processive exoribonucleases DIS3, DIS3L, DIS3L2, EXOSC10, XRN1, and XRN2. We initially optimized a competitive growth assay with engineered human Flp-In 293 T-REx cell lines that, upon induction with tetracycline, simultaneously express a short-hairpin RNA (shRNA) that silences an endogenous RNase gene and express an exogenous allele of that gene that is insensitive to the shRNA. Thus, we could replace the endogenous protein with a specific version that we chose. The construct also encodes a fluorescent protein that localizes to the nucleus, which serves as an shRNA expression marker (Fig. 1A). We established two stable cell lines for every RNase, in which the endogenous protein was replaced with an exogenous wildtype (WT) or mutated (MUT) version, and that simultaneously express enhanced green fluorescent protein (EGFP) or mCherry, respectively, so they can be cultured together and easily distinguished by fluorescence microscopy. The mutations we introduced completely inactivated DIS3L, DIS3L2, EXOSC10, and XRN2. In the case of DIS3, we introduced an MM-associated mutation that impairs exoribonucleolytic activity but does not destroy it completely because complete destruction would lead to cell death. Unfortunately, we were unsuccessful in generating cell lines with the inducible dysfunction of XRN1; therefore, this ribonuclease was not further investigated in the present study.

**Figure 1.**
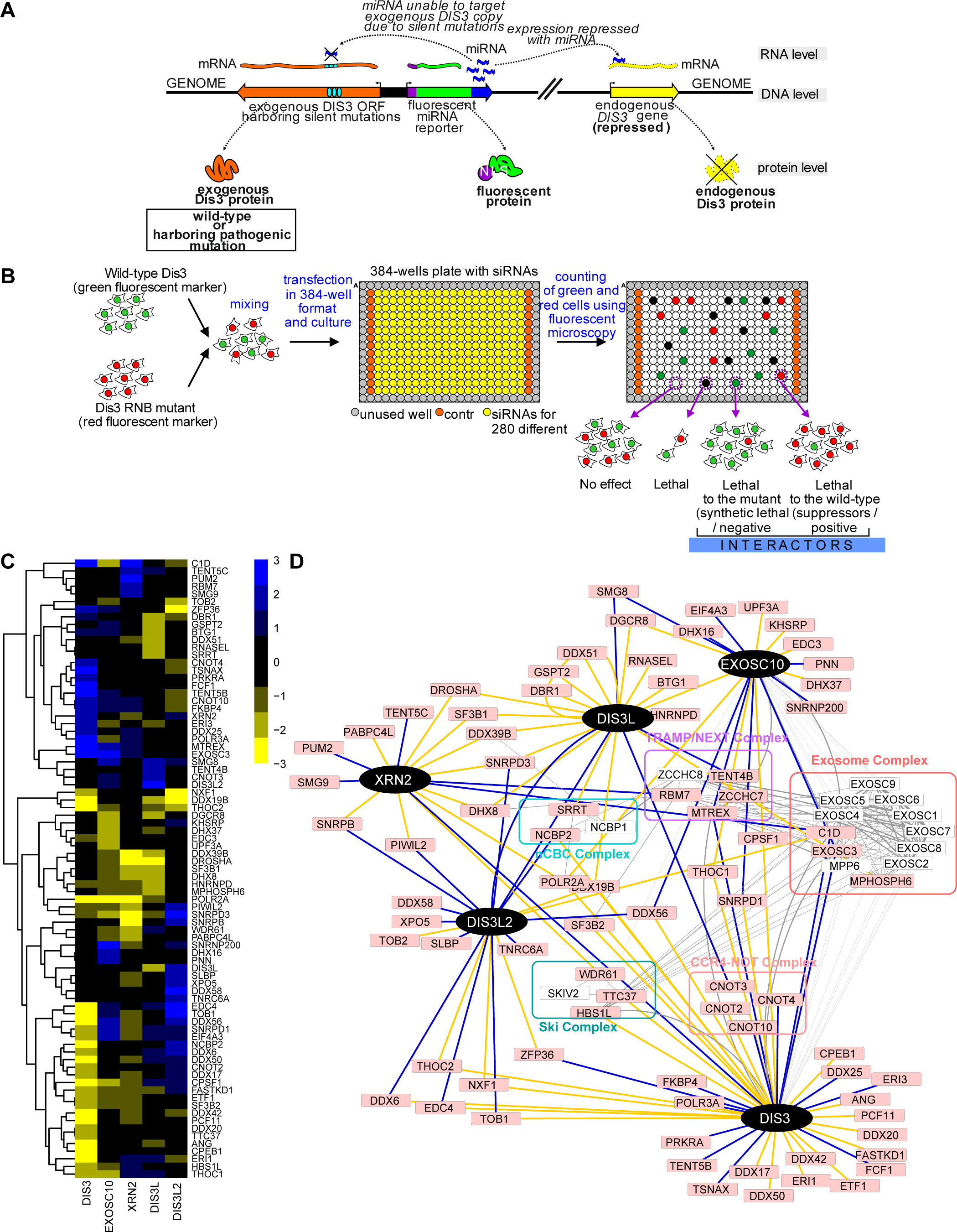
siRNA screens identified synthetic lethal interactions and suppressors of main mammalian processive ribonucleases. A. Strategy to obtain a cellular model in which the endogenous copy of the analyzed gene (DIS3 – as an example, DIS3L, DIS3L2, XRN2, or EXOSC10) is replaced by a mutant version. Schematics of a HEK293 Flp-In T-Rex cell line-based model are shown, in which a vector was integrated that contained a bidirectional promoter that – upon tetracycline-mediated induction – drove expression of shRNA that silenced the endogenous version of the gene and WT or MUT sh-miRNA-insensitive exogenous gene variant. B. Schematic presentation of the screening procedure. HEK293 Flp-In T-Rex cells that expressed WT and MUT versions of the analyzed ribonuclease were co-cultured in 384-well plates and transfected in triplicate with a custom or genome-wide siRNA library. The number of green (WT) and red (MUT) cells was quantified by fluorescence microscopy. Subsequently, a custom computational workflow was used to calculate fitness scoring based on changes in red and green cell count balance. Two types of GIs were identified: (*i*) negative/exacerbating (synthetic lethal), and (*ii*) positive/alleviating (suppressors). C. Heatmap of genetic interactionsof all of the analyzed ribonucleases, represented by summarized z* scores. The plots represent summarized z* scores for 81 of 280 tested genes that were identified as a significant hit for at least one query gene. Yellow represents positive hits (suppressors). Blue represents negative hits (synthetic lethal). Black represents genes that had no interactions. D. Network diagram of identified genetic interactions between mammalian processive ribonucleases and known physical interactions. Black nodes indicate query ribonucleases. Pink nodes indicate genes that were identified as hits in the siRNA screen. White nodes indicate proteins that interacted with query proteins or screen hits according to the STRING database. Blue and yellow edges represent synthetic lethal or suppressor interactions, respectively. Gray edges represent known P-P interactions. Protein complexes that are known to interact with query ribonucleases are highlighted in colors on the net.

All of the cell lines were interrogated with a library of siRNAs that targeted 280 preselected genes (Fig. 1B) that are involved in various aspects of RNA metabolism in the nucleus or cytoplasm (e.g., polyadenylation, transcription, translation, nuclear import/export factors, and some confirmed or predicted nucleases; Dataset EV1). The number of green (WT) and red (MUT) cells was measured by fluorescence microscopy 72 h after siRNA transfection (Fig 1B). We used custom computational workflow to calculate fitness scoring based on changes in red and green fluorescence balance and taking into account overall viability of cells in a given well (for details, see Materials and Methods). Thus, we were able to express the relationship between the fitness of WT and MUT protein-expressing cells as robust z scores (z* scores). Based on an arbitrary threshold of z* scores, two types of GIs were identified: (*i*) negative/exacerbating (synthetic lethal), in which the siRNA-mediated silencing of a target gene resulted in the specific growth inhibition or death of cells that expressed MUT protein, thus favoring the survival of cells with WT protein and (*ii*) positive/alleviating (suppressors), in which the siRNA-mediated silencing of a target gene resulted in the specific growth inhibition or death of cells with WT protein, thus favoring the survival of cells that expressed MUT protein.

After screening all of the processive nucleases, we found 111 strong GIs that involved 81 of 280 target genes in the library (Fig. 1C, Dataset EV2). Among the identified GIs we found well-established physical interactors of the analyzed ribonucleases (Fig. 1D). As expected for EXOSC10 and DIS3, a component of the exosome core (EXOSC3) was identified as a negative GI (Fig. 1D). C1D, a protein that is well known to interact with the exosome (Costello *et al*, 2011; Garland *et al*, 2013; Schilders *et al*, 2007), was classified as a hit for DIS3 and EXOSC10. Furthermore, components of protein complexes that are known to work as exosome activators were also identified as hits. MTR4, an RNA helicase that is encoded by the gene *SKIV2L2* gene, is part of the nuclear-localized Trf4/Air2/Mtr4p Polyadenylation (TRAMP) and nuclear exosome targeting (NEXT) complexes (Lubas *et al*, 2011; Puno & Lima, 2018; Weick *et al*, 2018; Meola *et al*, 2016), was also identified as a negative interactor with both DIS3 and EXOSC10. These results validated our screening methodology and allowed us to further evaluate with confidence previously unknown GIs of the main mammalian processive ribonucleases.

Interestingly, two superkiller (SKI) complex components (TTC37 and HBS1L), were found to be DIS3 mutation suppressors (Fig. 1D). The SKI complex is a major cytoplasmic exosome cofactor. It is responsible for cytoplasmic mRNA decay, in which it recruits NMD and non-stop decay (NSD) to the cytoplasmic exosome where they are degraded by DIS3L (Schmidt *et al*, 2016; Halbach *et al*, 2013). The reduction of cell viability that is caused by impairments in nuclear RNA degradation can be suppressed by impairing the cytoplasmic RNA decay process, indicating that the investigated genes function in pathways that are alternative ways with compensation result. Many genes that are involved in such suppressive interactions have close functional relationships(Leeuwen *et al*, 2016). A suppressor that functions in a related but alternative pathway and exhibits alterations of activity can restore or compensate for the dysfunction of a query gene by recreating conditions that are required for optimal cell growth. Similar to DIS3-SKI interactions, we found another example of the positive interplay between cytoplasmic and nuclear RNA decay. We found that the depletion of WDR61, another component of the SKI complex, has a suppressive effect on mutations of XRN2, another major ribonuclease in the nucleus. This complex network of interactions implies that nuclear degradation may somehow counterbalance cytoplasmic RNA degradation pathways.

In addition, among XRN2 negative hits were two more proteins that are known as exosome cofactors: C1D that is mentioned above and RBM7, a part of the NEXTcomplex. Intriguingly, those genetic interactions may suggest that in some context, XRN2 and nuclear exosome may work redundantly.

### Limited overlap between genetic interactions of the main RNA degrading enzymes

Several genes that are known to interact with the analyzed ribonucleases were identified as hits in our screens, thereby validating our experimental approach. To gain further insights into the genetic interaction network, we compared GIs for all five of the analyzed ribonucleases (Fig. EV1). Notably, we found that they tended to have nonoverlapping GIs, suggesting distinct properties that are associated with their loss of function. We did not identify any gene that showed a genetic interaction with all five enzymes. However, a few genes interacted with three ribonucleases. We identified five genes (6% of 81 hits) that had overlapping GIs with three ribonucleases (Fig. EV1 A and B). One apparent hit was the alpha subunit of RNAPII (POLR2A), which had a strong positive GI with all nuclear ribonucleases (DIS3, EXOSC10, and XRN2), suggesting that the inhibition of transcription can counterbalance dysfunctional RNA decay. The effect appeared to be more pronounced for nuclear RNases because cytoplasmic DIS3L and DIS3L2 did not interact with POLR2A, although such interactions were not studied for XRN1.

Another very strong positive hit was DDX19B, which was a suppressor of DIS3, DIS3L, and DIS3L2 (Fig. EV1 B). DDX19B is a highly conserved DEAD-box adenosine triphosphatase (ATPase) that is orthologous to yeast Dbp5p, an essential component of mRNA export machinery. Interestingly, Dbp5p works as an RNPase, rather than as an RNA helicase because it promotes the dissociation of Mex67p from mRNPs at the nuclear periphery (Lund & Guthrie, 2005; Tran *et al*, 2007). Consequently, disruption of the nuclear export process tends to have a suppressive effect on exosome-mediated alterations of RNA degradation. Intriguingly, the depletion of DDX19B can alleviate degradation defects that are mediated by all three DIS3 human paralogs, implying that paralog cooperativity might exist in some contexts.

Three additional genes, C1D, DDX56, and SNRPD3 were identified as interactors three times in our screen (Fig. EV1 B, D and E). But these interactions were much more complex, as they did not show a strict tendency to be synthetically lethal or suppressing with a specific subset of ribonucleases. For example, SNRPD3 was synthetically lethal with mutated DIS3L but was a suppressor of mutations of EXOSC10 and XRN2.

Twenty genes had GIs with two nucleases, and the other 69% of the genes that were identified as hits in our screens were unique GIs (Fig. EV1 A and C). Such a high number of uncommon GIs could be explained by the divergence of degradation processes that are mediated by those different ribonucleases.

From the perspective of interrogated ribonucleases, the highest number of common interactors (eight) was observed between DIS3 and EXOSC10. This was unsurprising because these nucleases share the same core complex, are present in closest cellular compartments, and have redundant functions in some contexts.

### Reconstruction of a functional network revealed novel connections between RNA metabolic pathways

To further reinforce the implications of the identified GIs with functions of each ribonuclease, we performed Gene Ontology (GO) enrichment analysis of genes that genetically interact with each of the query ribonucleases focusing mainly on negative interactions, as they tend to link genes in compensatory pathways.

The GO analysis of DIS3 negative GIs significantly showed enrichment of the rRNA processing term (Fig. 2A, and Dataset EV3), which is consistent with the description of the individual GIs that are mentioned above. Consistent with the cellular function of DIS3, two other biological processes that are related to rRNA processing were enriched: exonucleolytic trimming to generate the mature 3’-end of 5.8S rRNA from the tricistronic rRNA transcript and the maturation of 5.8S rRNA. Another significantly enriched term was nuclear-transcribed mRNA poly(A) tail shortening. Interestingly, genes that are ascribed to this biological process are a part of the CCR4-NOT complex (CNOT10 and CNOT4), and the third one (ZFP36) plays a role in recruiting CCR4 to this complex. CCR4-NOT is the major deadenylase that rapidly removes poly(A) tails from mRNAs that are to be degraded (Łabno *et al*, 2016; Azzouz *et al*, 2009). The exosome with DIS3 is the effector of this degradation pathway, so this GO term enrichment is consistent with established functional interactions of DIS3. Moreover, one of the GO terms that was enriched with negative interactors of DIS3 was gene silencing by RNA, which was enriched by three genes: TSNAX, CNOT10 and PRKRA. This was also consistent with known cellular functions of the exosome, which degrades the 5’ moiety of mRNAs that are cleaved by the RNA-induced silencing (RISC) complex during RNA interference in plants (Huntzinger & Izaurralde, 2011).

**Figure 2.**
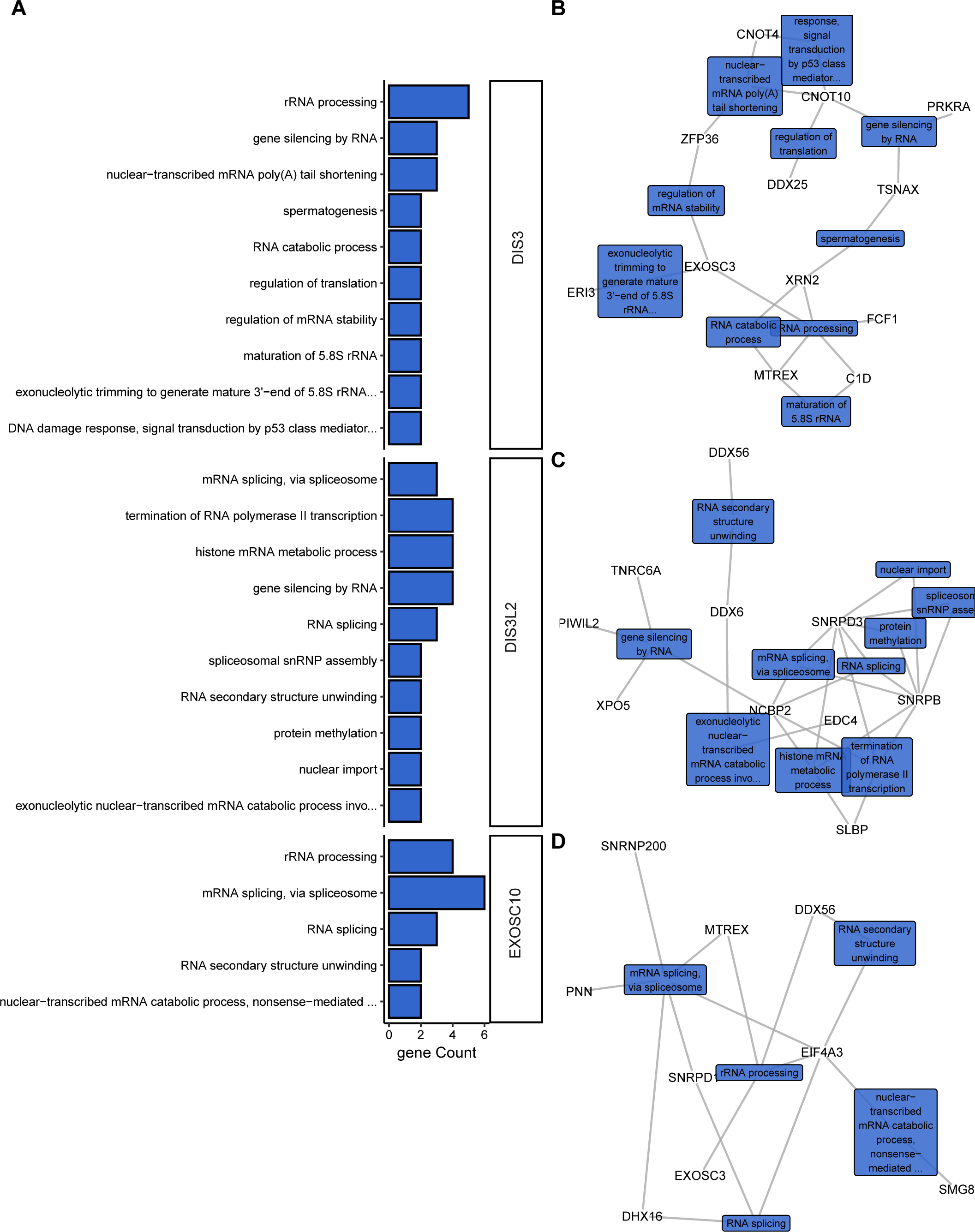
Functional analysis of synthetic lethal interactions revealed novel connections of mammalian ribonucleases with RNA metabolic pathways. A. Biological processes associated with genes that are synthetically lethal with mutations in DIS3, DIS3L2 and EXOSC10. Bar plots represent the number of identified negative hits associated with biological processes that were analyzed with the DAVID direct GO database. B-D. Functional interaction networks of DIS3 (B), DIS3L2 (C), and EXOSC10 (D) synthetic lethal interactors. Networks were constructed in R/Bioconductor based on the association of hit genes with biological processes according to the DAVID direct GO database. The colored rectangular labels show names of biological processes that are connected with gray edges to genes that are associated with them. No bar plots or network diagrams are shown for EXOSC10 and XRN2 because enrichment analyses of biological processes were non-significant.

Likewise, the GO enrichment analysis of negative interactors of EXOSC10 indicated that they are largely (four of nine) involved in rRNA processing (Fig. 2A, and Dataset EV3), which is consistent with the primary function of EXOSC10 in mammalian cells because it mainly resides in the nucleolus where the maturation of rRNA molecules occurs. Interestingly, the most enriched term was mRNA splicing via spliceosome, with six genes that were annotated to it: SNRNP200, EIF4A3, PNN, SNRPD1, SKIV2L2, and DDX16. EXOSC10 in budding yeast retains and degrades defective transcripts in the nucleus (Hilleren *et al*, 2001). However, the role of EXOSC10 in the surveillance of mRNA in mammalian cells has not been fully established. This suggests that EXOSC10 may play a role in cotranscriptional quality control in human cells, much like in yeast cells.

The GO enrichment analysis of negative interactors of DIS3L did not yield any significant terms because there were only four genes that were identified, and all of them are connected with RNA degradation. Likewise, GO enrichment analysis of XRN2 negative interactions did not yield any enriched Biological Processes.

Concerning negative genetic interactions of DIS3L2, three biological processes were significantly enriched: histone mRNA metabolic process, termination of RNA polymerase II transcription, and gene silencing by RNA (Fig. 2A and Dataset EV3). The enrichment of gene silencing by RNA and histone mRNA metabolic processes is consistent with known functions of DIS3L2 (Mullen & Marzluff, 2008). Enrichment of the termination of RNA polymerase II transcription term initially appeared surprising because DIS3L2 is a cytoplasmic protein, but genes that are ascribed to this process are also associated with the histone mRNA metabolic process term (Fig. 2C). Notably, these same genes are also connected with terms that are related to RNA splicing and mRNA export from the nucleus. These findings indicate that genes that are synthetic lethal with mutated DIS3L2 are involved in processes that are highly connected. Thus, speculating about which of these processes rely heavily on DIS3L2 to create strong genetic interactions that were observed is difficult.

### Distinct functional interactions were specific to nuclear and cytoplasmic exosome

DIS3 and DIS3L are close paralogs that have partially similar enzymatic activity and share the same core complex that is essential for their activity. Therefore, their functions may be at least partially redundant. However, they have different localization between the nucleus and cytoplasm, respectively. Such compartmentalization limits substrate availability and may also determine different interactions with biological processes. DIS3 and DIS3L likely perform divergent functions in the cell, which is consistent with our preliminary screens that revealed almost no overlap in GIs between DIS3 and DIS3L. To elucidate this issue and determine the biological processes that are affected by the loss of function of these two genes, we again performed a high-throughput siRNA screen using the same system but expanded the scope of the screen. We used a custom set of 3904 siRNAs that targeted genes whose products were directly or indirectly related to RNA metabolism (see Materials and Methods section for details, and Dataset EV1).

We identified 233 and 137 GIs that were specific to DIS3 and DIS3L, respectively, demonstrating that these nucleases play largely distinct roles (Fig. EV2). Only 15 genes interacted negatively with both DIS3 and DIS3L, indicating that although there might be some functional overlap, some processes in which these enzymes work cooperatively or redundantly are exceptions and not the rule. Moreover, 25 interactions were common for both nucleases but were suppressors of DIS3 mutation and synthetic lethal with MUT DIS3L. The complete list of GIs is provided in the Dataset EV4.

We analyzed 66 negative GIs that were specific to DIS3 with regard to biological processes but with hardly any meaningful enrichment. First, the enriched GO terms were represented by very few genes with no statistical significance (Fig. 3A and Dataset EV5). Second, these terms were not connected with one another, and the relationship between them and DIS3 is unclear (Fig. 3A). A small set of genes that are implicated in rRNA processing (EXOSC6, PES1, NCL, RPL6, and DIMT1) are the only exception. The limited enrichment in other than rRNA processing processes or pathways and a low number of negative GIs for DIS3 indicate that this enzyme is indispensable in its role. The importance of DIS3 as one of the main RNA degrading enzymes is confirmed by the fact that *DIS3* is an essential gene in all model systems that have been studied to date and indicates that its activity cannot be replaced even by the other nuclear exosome catalytic subunit, EXOSC10.

**Figure 3.**
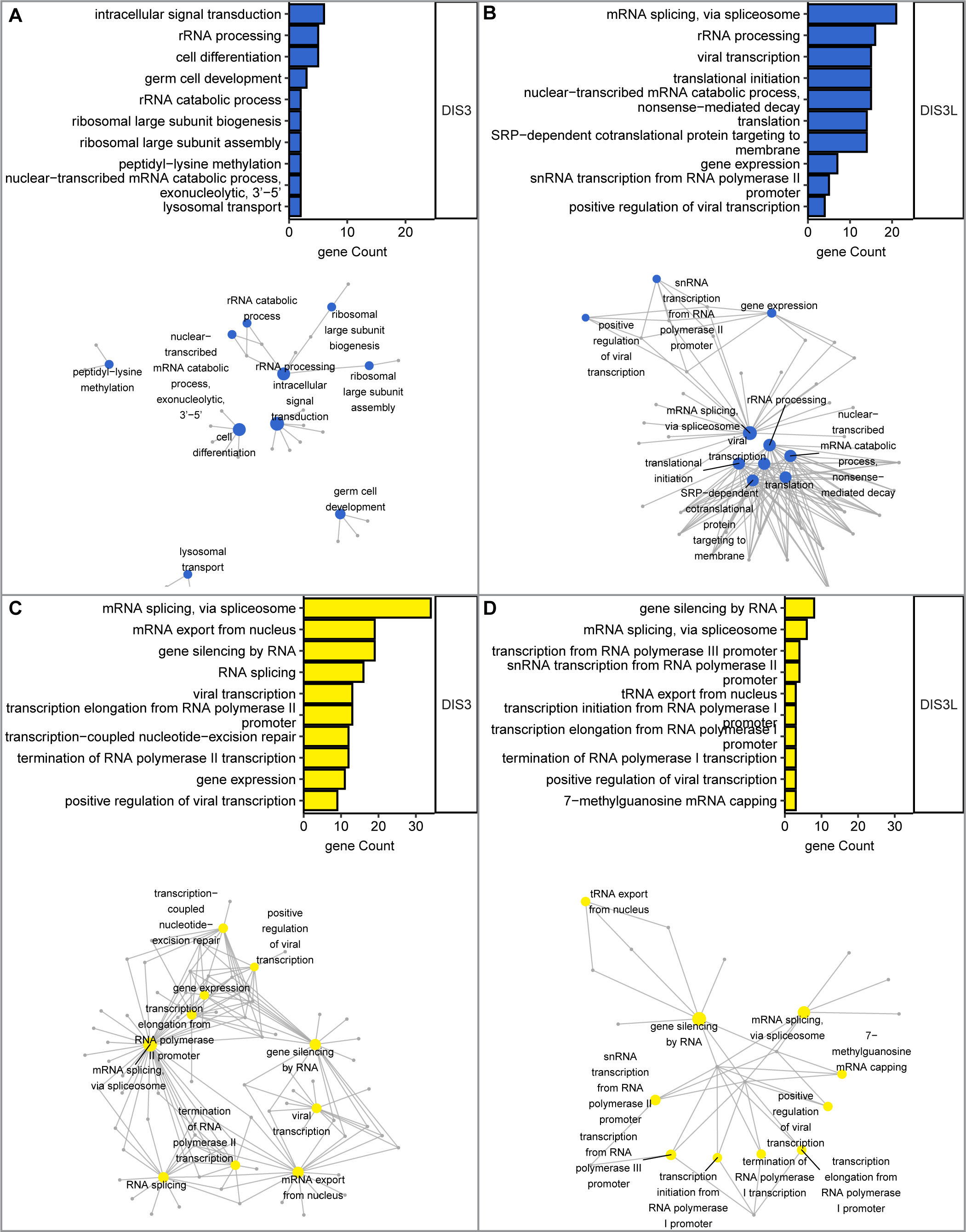
Distinct functional crosstalk between the nuclear or cytoplasmic exosome and RNA metabolism. A-D. Functional classification of synthetic lethal (A and B) and suppressor (C and D) interactions between DIS3 (A and C) or DIS3L (B and D) and RNA metabolism genes. Bar plots represent enrichment (expressed as gene count) among the top 10 biological processes (ordered by Fischer’s exact *p* value). The enrichment analysis was performed based on biological processes in the DAVID direct GO database. Networks illustrate connections between biological ontologies. The colored nodes (blue and yellow) represent biological processes that are interconnected with genes associated with them in GO. The networks were constructed in R/Bioconductor based on the association of hit genes with biological processes according to the DAVID direct GO database.

In contrast, the negative GIs of DIS3L fall into very specific GO categories (Fig. 3B). mRNA splicing and translation were among the highly enriched terms. The cytoplasmic exosome complex is involved in translation-dependent RNA surveillance pathways (Tuck *et al*, 2020; Schaeffer *et al*, 2011). Therefore, the enrichment of these GO terms corroborates its functions in RNA metabolism. Most of the enriched genes that are implicated in the translation process are ribosome complex constituents, from both small and large ribosome subunits. Genes that are enriched in the splicing category are constituents of the spliceosome from the second step of splicing and the U4/U6 x U5 snRNP complex. A high number of genes that are involved in splicing and were identified as synthetic lethal with MUT DIS3L suggests that the cytoplasmic degradation pathway by the exosome may somehow compensate for alterations of the mRNA maturation process. Notably, among negative GIs of DIS3L were genes that are involved in RNA transport from the nucleus (THOC2, ALYREF, NUP133, POP1, POP5, and EIF2S2). THOC2 and ALYREF are a part of the TREX complex (Chi *et al*, 2013), whereas NUP133 is a part of the nuclear pore (Boehmer *et al*, 2008), and all are involved in mRNP biogenesis and mRNA export (Vasu *et al*, 2001; Fan *et al*, 2019). These steps are subject to robust quality control. These results affirm the possibility that cytoplasmic degradation may compensate for the deregulation of nuclear mRNA metabolism.

Among the positive GIs were 15 that were common between DIS3 and DIS3L. Within this group, we found two interesting categories: (*i*) genes that are involved in RNA export from the nucleus that are part of the nuclear pore complex (RAN, TPR and NUP153) and (*ii*) subunits of the RNA polymerase complex (POLR1C, POLR2G, POLR2H, and POLR2L).

Both paralogs interacted with RNA polymerases I and II, but DIS3 showed a more potent genetic interaction with RNA synthesis machinery, as almost all subunits of RNAPII (POLR2A, POLR2B, POLR2D, POLR2H, POLR2G, POLR2F, POLR2L, and POLR2I), one subunit of RNAPI (POLR1C), and one subunit of RNAPIII (POLR3K) were identified as positive GIs.

Remarkably, positive GIs that are specific to DIS3 fall into the most diverse GO categories. The highest number of genes were enriched in categories that are connected to mRNA splicing (Fig. 3C). This result is unlike the findings for DIS3L. For this nuclease, more GIs that are implicated in the splicing process were negative GIs, however, among suppressors the mRNA splicing was also enriched (Fig. 3D). Markedly, distinct sets of genes were enriched in the splicing process for DIS3 and DIS3L. One commonality was that most of them were constituents of the catalytic step 2 spliceosome. The second process that was the most enriched was mRNA export from the nucleus. Within this set of genes were parts of the nuclear pore and nuclear envelope.

Altogether, these results suggest that lower RNA synthesis or RNA export from the nucleus to the cytoplasm counteracts the dysfunction of exosome-dependent RNA decay.

### DIS3 dysfunction suppressed alterations of alternative splicing in cells with the depletion of U2 snRNP component

Considering the above observations, we were intrigued that genes that are implicated in splicing were the most abundant ones among MUT DIS3 suppressors. Moreover, the fact that constituents of U2 snRNP and U5 snRNP were the most enriched group among positive hits suggests that DIS3 dysfunction leads to alterations of splice site recognition. These results raised the possibility that the interplay is between alterations of splicing and the dysfunction of DIS3. Thus, we dissected the effect of the depletion of one of the strongest positive hits. Among these were components of the splicing factor 3A subunit 1 (SF3a) complex and SF3b complex. The SF3a complex is an integral part of the pre-spliceosome and is required for complex A and B assembly (Brosi *et al*, 1993). SF3a interacts with U1 and U2 snoRNAs and pre-mRNA and plays a crucial role in initial steps of spliceosome assembly (Lardelli *et al*, 2010; Bertram *et al*, 2017; Tanackovic & Krämer, 2005; Fica & Nagai, 2017). We used one of its subunits (SF3A1) to study the interplay between the exosome and splicing. We examined the way in which the dysfunction of DIS3 affects splicing after SF3A1 depletion.

To profile changes of alternative splicing (AS) patterns, we performed total RNA sequencing (RNA-Seq) under four conditions: cells that expressed either DIS3^WT^ or the mutated version DIS3^G766R^ that were treated with either siRNA targeting the SF3A1 gene (siSF3A1) or control siRNA (siCTRL). To compare profiles of AS events, we employed JUM pipeline (Wang & Rio, 2018) because it allows the identification and estimation of global splicing patterns and differential analysis by combining many experimental conditions and considering biological replicates. JUM handles RNA-Seq reads that are mapped to splice junctions and first defines alternatively spliced junctions, assembles them into common known AS events (alternative 5’ splice site [A5SS], alternative 3’ splice site [A3SS], cassette exon [SE], mutually exclusive exon [MXE], and intron retention [IR]), and creates an additional separate category for complex AS isoforms (MIX). Expression levels of each identified AS is quantified by computing the change in percent spliced isoform (ΔPSI). Differential inclusion is usually restricted for |ΔPSI|> 0.1. Significantly different AS events can be identified by applying statistical tests (see Materials and Methods). Based on this approach, we were able to identify condition-specific splice junction usage and check whether there are any significant differences in AS event frequencies between conditions.

The analysis revealed that SF3A1 depletion led to extensive alterations of multiple classes of AS events (Fig. 4A). Importantly, significantly deferentially spliced events were not distributed equally between classes: SE was notable with 460events compared with 180 events for IR, 151 events for A3SS, 147 events for A5SS, and 106 events for other classes.

**Figure 4.**
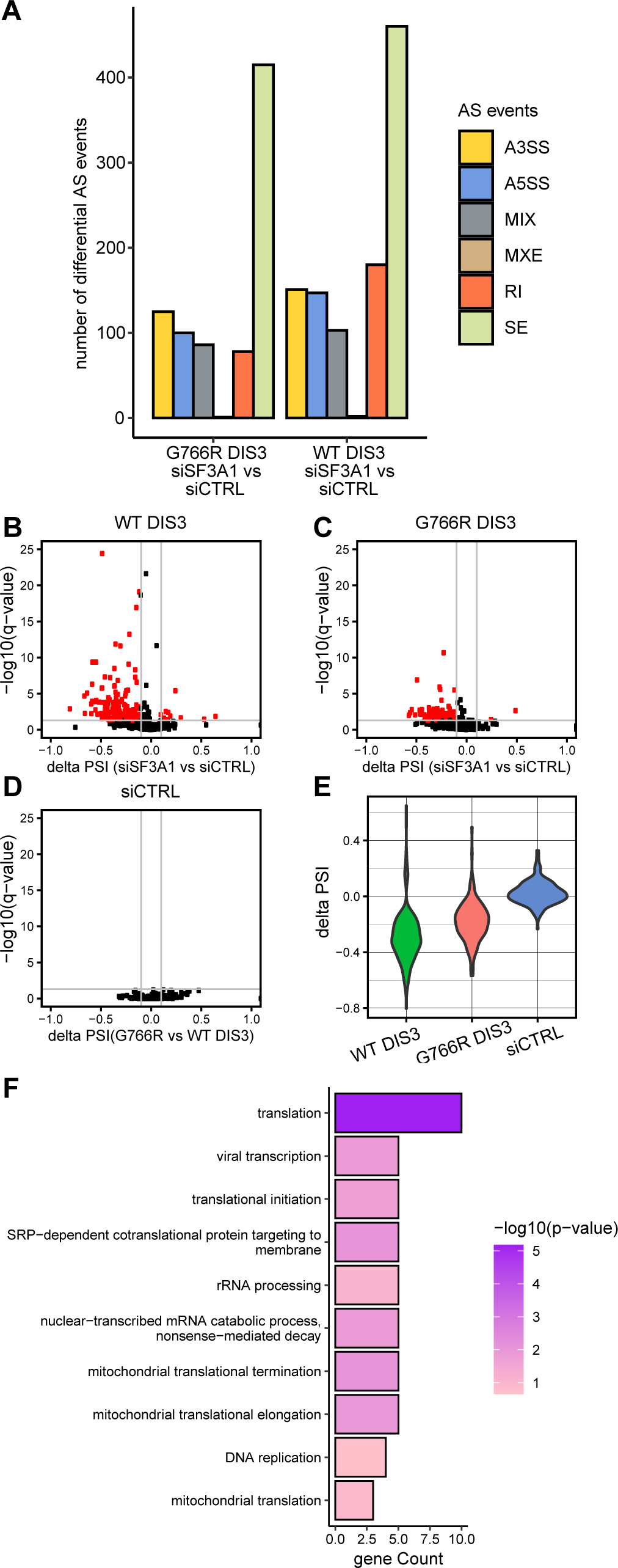
Suppression of alterations of alternative splicing that are caused by the depletion of a U2 snRNP component in the presence of DIS3 dysfunction. A. Quantitative comparison of AS events with significant alterations of splicing patterns after SF3A1 depletion in cells that expressed DIS3^WT^ or DIS3^G766R^ (deferentially spliced events with |ΔPSI| > 0.1 and FDR < 0.05). Analyzed AS events: alternative 5’ splice site (A5SS), alternative 3’ splice site (A3SS), cassette exon (SE), mutually exclusive exon (MXE), intron retention (IR), and complex AS isoforms (MIX). B-D. Volcano plots representing the direction and significance of retained intron changes, analyzed with JUM. Every dot is an identified IR event. Red dots are IR events that were significantly changed between SF3A1 depletion and control conditions (|ΔPSI| > 0.1 and FDR < 0.05). E. ΔPSI violin plots of the distribution of calculated retained isoforms of introns that were significantly alternatively spliced under SF3A1 depletion conditions in a DIS3^WT^ background across other analyzed conditions (*i.e.*, in cells that expressed DIS3^G766R^ and when comparing splicing between DIS3^WT^ and DIS3^G766R^ in siRNA control transfection [siCTRL] conditions). F. Gene Ontology enrichment analysis of biological process terms of JUM-identified transcripts with retained introns after SF3A1 depletion. Bar plots represent the number of identified genes that were associated with biological processes, analyzed with the DAVID direct GO database. The bar color code shows statistical significance, colored according to the value of negatively log10-transformed Fischer’s exact *p* value.

Remarkably, DIS3 dysfunction strongly altered the effect of SF3A1 silencing on only two classes of AS events: IR and A5SS. Among 591 of the identified IR events, SF3A1 depletion in the presence of DIS3^WT^ significantly altered 180 introns, and 169 of these changes were more retained introns, meaning that retention frequency was higher than under control conditions (DIS3^WT^ cells treated with siCTRL) (Fig. 4B). In the presence of DIS3^G766R^, only 78 IR events were significantly altered (Fig. 4C), whereas mutation in DIS3 alone had no influence on intron retention (Fig 4D). Notably, for some of the introns SF3A1 depletion caused retention with a ΔPSI < −0.5 and median around −0.35 in the presence of DIS3^WT^ (Fig. 4E). In the presence of DIS3^G766R^, the median ΔPSI was not below −0.25. More detailed analysis revealed that more than 30% of retained introns were the first intron of the transcript. Further evaluation, however, did not show any correlation between the presence of these introns in the 5’ untranslated region (UTR) or coding sequence (CDS). To investigate the biological processes in which the transcripts that retained their introns may be implicated, we performed GO enrichment analysis. Interestingly, translation was the most enriched GO term (Fig. 4F). The other significantly enriched terms were also related to translation. The precise examination of genes that were enriched in these GO terms revealed that they encode ribosomal proteins (RPs): RPL10, RPL10A, RPL24, RPL41, RPL27A, MRPL52, MRPS12, MRPS24, and MRPS34.

This suggests that DIS3 dysfunction alleviates SF3A1 depletion by rescuing appropriate isoforms of transcripts that encode ribosomal proteins, which in turn may influence translation and amend the balance between RNA processing, RNA degradation, and translation and help maintain cell homeostasis.

Several conclusions emerged from this experiment. First, SF3A1 depletion does not dysregulate splicing globally but rather leads to specific AS alterations, which is consistent with previous reports. The knockdown of individual splicing factors has noticeable but very distinct effects on AS (Papasaikas *et al*, 2015). Second, DIS3 dysfunction suppresses alterations of splice site selection upon SF3A1 depletion but only in IR and A5SS classes of alternative splicing, suggesting a particular mechanism of action rather than a global one. Third, in SF3A1-undepleted cells, the AS pattern was not significantly altered in DIS3^G766R^ compared with DIS3^WT^, and we did not observe the accumulation of specific pre-mRNAs.

In conclusion, our data indicate that DIS3 dysfunction alleviates the effects of SF3A1 depletion on a very specific subset of genes, which partially explains the suppressor interaction that was observed in the genetic screen. Further investigations are necessary to decipher the molecular mechanisms that determine why this particular subset of genes is a target of the DIS3-SF3A1 interaction.

### Genome-wide siRNA screen of DIS3 identified novel interactions of the nuclear exosome with processes that are involved in maintaining cell homeostasis

The fact that DIS3 is linked to human disease and appeared to be engaged in an interplay with different cellular pathways prompted us to expand our siRNA screen to the whole-genome scale. We performed a genome-wide siRNA screen using a commercially available siRNA library that covers most human genes (Dharmacon ONTARGETplussiRNA SMARTpool; 18104 targets).

Expanding the screen form 3904 genes that are involved in RNA metabolism to nearly the entire genome increased the number of observed interactions from 274 to 1112. This number, although substantial, is lower than could have been expected given the size of the library. The complete list of GIs is provided in the Dataset EV6.

Functional analysis of the entire collection of genes that genetically interact with DIS3 revealed that the most enriched categories are consistent with the analysis of GIs that was performed with the custom RNA metabolism library (Fig. 5A and B, and Dataset EV7). Notably, the functional categories that were most extensive and represented at significantly higher frequencies than expected by chance were again connected with mRNA splicing, the regulation of RNAPII transcription, and mRNA export from the nucleus (Fig. 5B). Nevertheless, we investigated whether genes that genetically interact with DIS3 are associated with biological processes other than RNA metabolism. We reevaluated the functional categorization of DIS3 GIs, excluding genes that were previously examined. The GO analysis of these positive and negative genetic interactors revealed that they fall into a very diverse group of functional categories that are important for cell maintenance (Fig. 5A and B, Fig. EV3). Importantly, the detailed examination of enriched categories revealed that they are enriched by a considerably low number of genes, meaning that genes are rather uniformly distributed among functional categories, with no category that is represented by a significant number of genes (Fig. 5A and B). Analyses of the relationships between these genes did not yield a highly connected interaction network, which was in contrast to GIs that were identified among genes that are involved in extended RNA metabolism Fig. EV3). Therefore, we subsequently performed an analysis of functional connections among enriched biological processes and cellular compartments. These analyses yielded some overrepresented functional categories that grouped together (Fig. EV4, EV5). These functional categories point to processes in which DIS3 may play an important role. We do not discuss all of these interactions in detail but highlight those that can provide essential information for further studies of the role of DIS3 in diverse biological processes.

**Figure 5.**
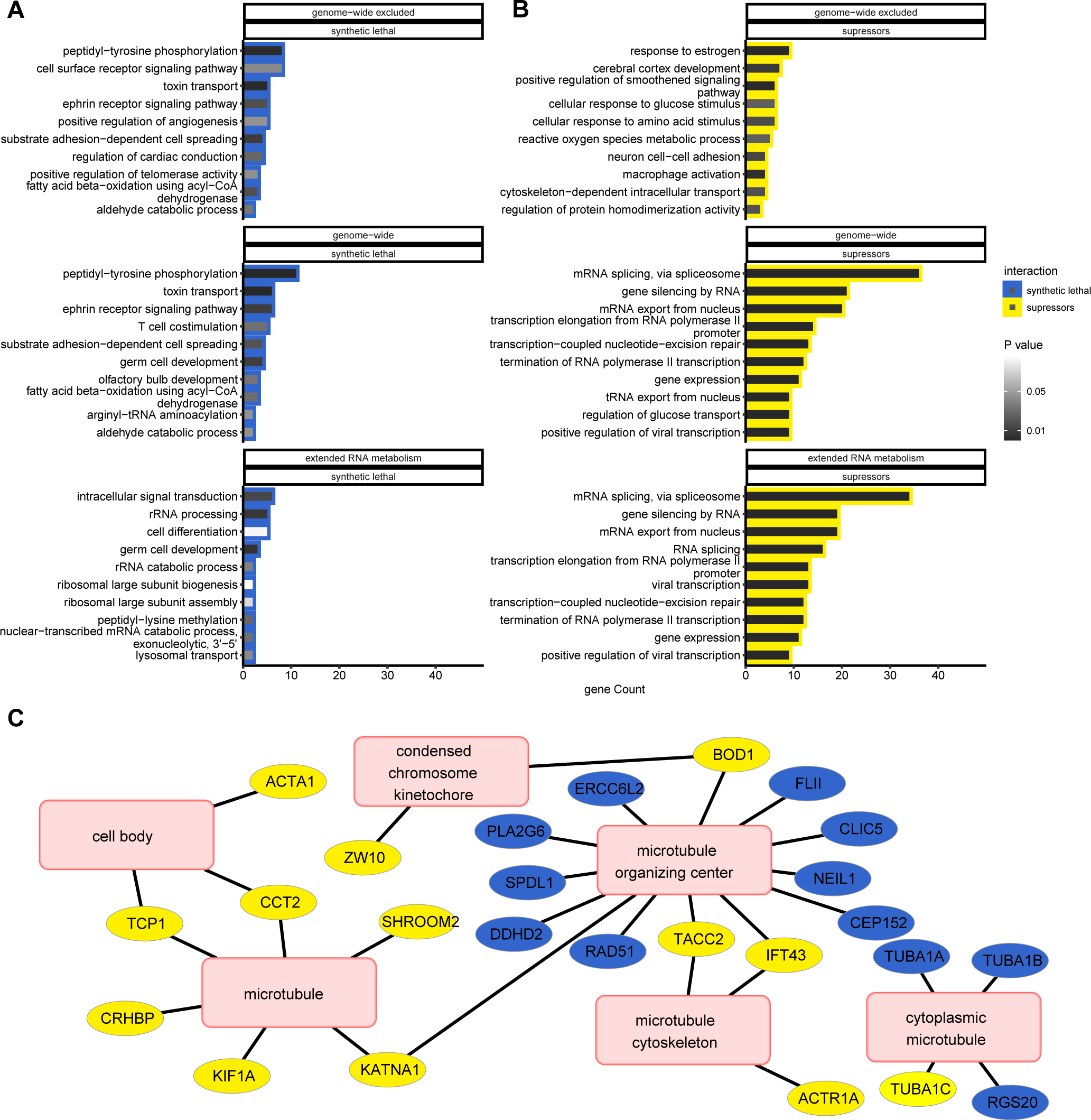
Genetic interactions between the nuclear exosome with genes that are involved in the maintenance of cell homeostasis. A-B. Comparisons of biological processes that were associated with genetic interactions of DIS3 identified in the genome-wide screen, extended RNA metabolism screen and hits form genome-wide subset, excluding hits that were identified in the extended RNA metabolism screen. Bar plots represent the number of identified hits that were associated with biological processes according to the enrichment analysis using the DAVID direct GO database. The fill color of the bar plots represents Fisher’s exact *p* value. Enrichment analyses were performed separately for (A) synthetic lethal interactions and (B) suppressors. C. Functional connections of synthetic lethal interactors and suppressors of DIS3 that are associated with microtubules. The hits of enriched cellular compartment categories that were connected to microtubules were extracted from the enrichment results using the DAVID direct GO database and visualized in Cytoscape as a network in which edges represent association of genes (blue nodes for synthetic lethal interactions and yellow nodes for suppressors) with functional categories (pink nodes).

We used DIS3 with a mutation in the RNB domain that was reported to be present in MM patients. We sought to identify new genetic interactions of DIS3 that could be relevant to MM. Of the genes that are reportedly involved in MM, only three had significant differential effects on cell viability in our screen: *IRF4*, *CCND1*, and *MAFB*. *IRF4* encodes a transcription factor that is involved in the interferon response. It is often upregulated in MM because of t(6;14) translocation and has been described as required for MM survival (Schaeffer *et al*, 2009). The silencing of this gene decreased the viability of cells that had the DIS3^G766R^ mutation. *CCND1* encodes a cyclin that is involved in G1/S progression, which is positively regulated by and interacts with RB1. It is frequently mutated in MM (Lawrence *et al*, 2013). The silencing of *CCND1* confers an advantage for cells with MUT DIS3, which corroborates data in the literature. *MAFB* encodes a transcription factor that is involved in hematopoiesis. It is upregulated in MM cases with t (14;20) translocation (Prideaux 2014). The silencing of this gene negatively affected all of the cells, and cells that expressed WT DIS3 were affected more severely.

Additionally, we looked deeper into genes that to date have not been associated with MM. Among negative GIs of DIS3, we found genes that are involved in the regulation of telomerase activity (CCT2, NVL and TCP1) (Fig. 5A and Dataset EV7). This was surprising because previous studies indicated that knockdown of the exosome rescued defects in telomerase activity and suggested that the exosome is a target for specific telomerase pathologies (Shukla *et al*, 2016). This inconsistency could be attributable to the fact that the exosome is involved in degradation of the telomerase RNA component (hTR) (Tseng *et al*, 2018, 2015; Gable *et al*, 2019; Nguyen *et al*, 2015), and rescue by exosome inactivation restored defective telomerase activity through lower levels of hTR (Boyraz *et al*, 2016). In contrast, genes that were identified in our screen participate in assembly of the telomerase holoenzyme. NVL can regulate telomerase assembly via its interaction with hTR (Her & Chung, 2012, 2), whereas CCT2 and TCP1 are part of the TRiC complex, which is required for TCAB1 protein folding (Freund *et al*, 2014). The depletion of TRiC subunits led to the mislocalization of hTR to nuclei and not to its global reduction (Chen *et al*, 2020, 1). Moreover, hTR overexpression cannot compensate for TRiC or TABC1 loss. Although this explains why DIS3 dysfunction did not suppress the loss of proteins that regulate telomerase activity, it does not explain the observed synthetic lethality. Thus, the role of DIS3 in maintaining telomerase activity appears to be much more complex and should be further investigated.

Further analyses of GIs that were identified in the genome-wide screen also revealed a substantial number of genes that are involved in microtubule organization (Fig. 5C, Dataset EV7). DIS3 dysfunction reduces cell growth and division. Cells that lack DIS3 activity exhibit the presence of overcondensed chromosomes (Snee *et al*, 2016) and are blocked in mitosis (Murakami *et al*, 2007). Moreover, DIS3 mutants in yeast are synthetically lethal, with mutations that influence the kinetochore, anaphase-promoting complex, or spindle pole body (Smith *et al*, 2011; Milbury *et al*, 2019). The results of our screen indicated that the interplay between DIS3 deactivation and microtubule organization is much more diverse. We observed genes that belong to microtubule or microtubule organizing center as both suppressors and synthetic lethal interactions. Comprehensive analyses of GIs that are engaged in microtubule organization showed some tendencies as synthetic lethal interactions include genes that are involved in the kinetochore and cell body, which is consistent with previous studies (Smith *et al*, 2011; Milbury *et al*, 2019). Positive interactors of DIS3 include genes that are involved in the motor function of microtubules and are also involved in the DNA damage response. This result provides a valuable resource to direct future studies of the involvement of DIS3 in microtubule maintenance.

## DISCUSSION

Understanding the effects of complex functional connections between gene products and their outgrowth on complex relationships between genotypes and phenotypes remains a fundamental challenge in modern genetics (Costanzo *et al*, 2019). In the present study, we applied high-throughput siRNA screening to map GIs of the main mammalian ribonucleases: DIS3, DIS3L, DIS3L2, EXOSC10, and XRN2. Using a combination of experimental and computational approaches, we obtained a dataset that allowed us to explore the ways in which key players in RNA decay collaborate with diverse cellular processes. Three main conclusions can be formulated from our results. First, both highly processive nuclear ribonucleases DIS3 and XRN2 are indispensable in their functions, whereas EXOSC10 and the cytoplasmic ribonucleases DIS3L and DIS3L2 appear to be auxiliary. Among the negative GIs of DIS3 and XRN2, there was no specific enrichment in particular biological processes. Second, the alleviating GIs of all of the query ribonucleases at the level of biological processes are highly connected. Moreover, most of the genes that were identified as positive hits are not directly involved in RNA degradation but rather other RNA metabolism processes, such as transcription, pre-mRNA splicing, and nuclear export. Cytoplasmic ribonucleases genetically interact with genes that are involved in nuclear processes. Third, DIS3 and DIS3L possess unique functions in RNA degradation. Moreover, the investigation of DIS3 and DIS3L interactomes showed that they interact in opposite ways with some processes, such as splicing. This is a good example of functional divergence, particularly subfunctionalization, in which both paralogs are separately specialized in several ancestral functions.

The GIs that were mapped in the present study support the unique functions of each of the examined ribonucleases, which is consistent with distinct substrate specificity (Szczepińska *et al*, 2015; Tomecki *et al*, 2010; Staals *et al*, 2010; Ustianenko *et al*, 2013; Davidson *et al*, 2012, 2019; Lubas *et al*, 2013), which can be explained by their different modes of substrate selection, catalytic activity, and cellular compartmentalization in mammalian cells. Interestingly, in our preliminary screens, we restricted the tested library to genes that are closely related to RNA metabolism. More than 65% of the mapped interactions were unique, indicating that these ribonucleases indeed operate separately. However, we also detected some shared GIs that may indicate that these enzymes either cooperate or are redundant in contexts of selected biological processes. Some of the overlapping GIs can be easily explained as negative interactions between EXOSC10 and DIS3 and the core exosome component EXOSC3 or exosome cofactors C1D and MTR4. One example of redundancy in utility is MTR4, which is known to play an essential role in all aspects of nuclear exosome functions by targeting various RNA substrates to it. In the nucleolus in mammalian cells, it associates with other proteins to form the TRAMP-like complex that is involved in rRNA processing, in which EXOSC10 plays a crucial role as the exosome’s executive nuclease. MTR4 is also present in the nucleoplasm where it forms the NEXT complex that is responsible for recruiting a broader group of exosome substrates, such as PROMPTs, snRNAs, and pre-mRNAs, which are degraded by DIS3.

The novel GIs that were identified in the present study are common to somewhat autonomous RNA degradation pathways and reveal novel aspects of functional links. The reconstruction of some of the known complexes on the GI networks indicated novel compensation or cooperation possibilities in crosstalk between RNA decay pathways. For example, among the genes that aggravated the XRN2 mutation, we identified genes that work as exosome cofactors (RBM7 and C1D), implying that the exosome may work redundantly with XRN2 in the nuclear degradation of some RNA species. XRN2, similar to the exosome, is involved in the rRNA maturation process by processing 5’ ends and removing excised spacers (Sloan *et al*, 2013; Preti *et al*, 2013; Wang & Pestov, 2011). The processing of 5’ends by XRN2 provides a quality check to decide whether the pre-rRNA should be further processed or degraded (Sloan *et al*, 2013; Wang & Pestov, 2011). A yeast homolog of C1D, Rrp74p, was shown to be required for exosome-mediated rRNA processing (Mitchell *et al*, 2003). This indicates the redundancy of 5’ and 3’ nuclear processing pathways for some stable RNAs is possible.

The structural and biochemical characterization of main executors of RNA processing, degradation, and turnover, including mechanistic details, were elucidated in detail, but their genetic interactions have not been systematically explored previously. To date, the only knowledge of genetic interactions has been available for yeast exosome-related ribonucleases (Dis3 and Rrp6) and can only be extracted from global genetic interaction networks for nonessential and essential budding yeast genes (Costanzo *et al*, 2016; Leeuwen *et al*, 2016; Usaj *et al*, 2017). The results from these previous studies partially overlap with the present results, in which they indicate that yeast Dis3 and Rrp6 are mainly involved in rRNA processing. These results are also consistent with our conclusion that DIS3 is required for its function.

During mapping GIs, more emphasis is usually given to GIs that more severely decrease cell fitness than to suppression interactions. The reason for this may be the difficulty in explaining the underlying mechanisms between these types of communications. However, Leeuwen et al. studied on the global scale genetic suppression interactions in budding yeast and classified them into mechanistic categories (Leeuwen *et al*, 2016). They showed that a substantial part of positive GIs can be explained by already known functional interactions, such as when query genes and suppressors are from the same complex or the same or alternative pathway. However, ∼50% of cases of suppression interactions are with unknown, apparently functional relationships, thus providing a good source of compensatory pathway identification. The analysis of GIs that were identified by us in biological processes revealed important interactions of RNA degradation in a broad biological context and uncovered a complex network of genetic suppression interactions. The analysis of positive GIs that were identified in our screens uncovered important reciprocal actions between RNA decay and transcription. For all of the analyzed nuclear ribonucleases, we observed the strong suppression of phenotypes that are connected with their alterations of activity when POLR2A was depleted. Moreover, at the biological process level, among the mapped GIs were genes that are involved in the process of transcription termination. The genome-wide DIS3 screen revealed that all RNAPII subunits and a broader group of genes that are involved in transcription regulation at the level of elongation and termination are positive interactors of this ribonuclease. The observed cooperation between RNA synthesis and the degradation process indicates the ways in which cell survival depends on coordinated actions between those two fundamental processes. It also implies an important role for proper buffering of the imbalance between these processes. The concept of maintaining a balance between RNA degradation and synthesis is not new, and some attempts have been made to investigate it in the past (Rogowska *et al*, 2005). However, most of them concern cytoplasmic mRNA degradation and its interplay with RNA synthesis (Sun *et al*, 2013; Shalem *et al*, 2011; Goler-Baron *et al*, 2008; Dori-Bachash *et al*, 2011). Our data also reveal an interplay between nuclear degradation processes and RNA synthesis. Moreover, our results clearly indicate that the deregulation of RNA synthesis may compensate alterations of RNA decay. Further work is necessary to shed light on the molecular mechanisms that underlie this interplay.

The evaluation of positive GIs also revealed the extensive interplay between RNA decay and RNA splicing. Surprisingly, the connection between these processes is mainly through alleviating interactions. This association was strongly highlighted by the results of the genome-wide screening of DIS3, in which splicing was the most prominent biological process that was enriched by genes that were identified as positive interactors. The cooperation between RNA splicing and exosome-mediated RNA degradation remains poorly understood. Our attempt to decipher the molecular mechanism of this interplay showed a very specific pool of transcripts where alterations of DIS3 activity mitigate the dysregulation of alternative splicing that is caused by the depletion of one splicing factors. Thus, the observed effects were not global. However, most of the transcripts that were rescued to the correct splicing isoform encode ribosomal proteins, and the observed effect of suppression on cell fitness was strictly connected with the restoration of appropriate levels of these mRNAs and their protein products.

Finally, the genome-wide DIS3 high-throughput siRNA screen allowed us to functionally characterize this ribonuclease and provide insights into processes other than RNA decay in which it participates. The results of this screen may also be a rich resource for other investigators because we only described a small number of possible interactions. The present findings may help direct further explorations of connections of DIS3 mutations to different phenotypes of cellular physiology.

In summary, the present data improve our understanding of the interplay between RNA decay and other cellular processes. To our knowledge, this was the first attempt to globally dissect genetic interaction networks that focus on RNA processing by employing large-scale analyses.

## MATERIALS AND METHODS

### Cellular model and cell line generation

Each screen was a competitive growth assay in which the fitness of cells that express the MUT protein of interest was compared with cells that expressed WT protein. We used an approach that was previously described by us to construct the cell lines (Tomecki *et al*, 2014; Szczepińska *et al*, 2015; Szczesny *et al*, 2018). Briefly, the parental 293 Flp-In T-REx cell line was stably transfected with a construct in which a bidirectional, tetracycline inducible promoter controls the expression of two transcription units. One transcription unit produces a bicistronic RNA that contains a miRNA precursor and mRNA for a fluorescent protein that localizes to the cell nucleus and serves as an expression marker. The other transcription unit contains the cDNA of the gene of interest with a C-terminal FLAG tag. The cDNA contains silent point mutations that render the resulting mRNA insensitive to the miRNA. Thus, the endogenous protein is replaced by a desired version of the protein that is encoded by the transgene and expressed at roughly the same levels. For every gene of interest, two stable cell lines were generated: one that expressed the WT version of the respective protein and EGFP as the expression marker and the other that expressed the protein of interest with mutations that impaired or abolished its catalytic activity with mCherry as the expression marker. This allowed the identification of cells that express the miRNA and distinction between WT and MUT cells.

### Cell culture and transfection

293 Flp-In T-REx cells were cultured in Dulbecco’s Modified Eagle Medium (DMEM) with 10% fetal bovine serum (FBS) without antibiotics at 37°C in a 5% CO_2_ atmosphere, with passages every 2-3 days. The cells that were used for transfection were obtained from 2-10 passages after thawing. Screening was performed in triplicate in 384-well plates (Greiner) with a poly-L-lysine-coated uClear bottom. Target siRNAs were stored at −80°C in 384-well plates. The day before transfection, they were thawed, and an aliquot was transferred to a master plate. Control siRNAs were added to the master plates by hand, and then the master plate was replicated to transfection plates. The plates were then sealed and stored at 4°C overnight. The next day, Lipofectamine siRNA MAX was added to the wells in of OptiMEM. This was performed with a MobiTek dispenser.

WT and MUT cells were grown separately. On the day of transfection, they were detached, counted, and mixed at a ratio where untreated cells reach roughly equal numbers (green:red) after 72 h of culture. This plating ratio was determined for each cell line and each batch of thawed cells. Once the cells were mixed, they were brought to an equal concentration, and 750 cells per well were added to the transfection plates with a MobiTek dispenser. The plates were resealed and left at room temperature for 1 h to allow the cells to evenly attach to the plate surface. Afterward, they were moved to an incubator and cultured for an additional 71 h.

Seventy-two hours after transfection, 10% formaldehyde in phosphate-buffered saline (PBS) with 2 mg/ml Hoechst 33342 was added to each well, and the cells were fixed and stained for 20 min at room temperature. They were then washed three times with PBS and covered with 50 µl of PBS with 0.1% sodium azide. The plates were sealed and stored at 4°C until imaging but for not more than one week.

### siRNA libraries

Human ON-TARGETplus SMARTpool siRNAs (Dharmacon) were used. A pool of four different siRNAs for each gene were used in a 384-well format. The core RNA metabolism library was designed by manually selecting 280 genes that are involved in various aspects of the RNA life cycle, including known and predicted RNA helicases that were described previously (Jankowsky *et al*, 2011). The extended RNA metabolism library was created by selecting 3904 siRNA pools from the Human Genome ON-TARGETplus siRNA Library-SMARTpool (Dharmacon). Genes that were targeted by the extended RNA metabolism library were chosen based on results of the experimental capture of RNA binding proteins (Castello *et al*, 2012; Beckmann *et al*, 2015; Baltz *et al*, 2012; Castello *et al*, 2016; Conrad *et al*, 2016; Kwon *et al*, 2013; He *et al*, 2016; Liao *et al*, 2016), the prediction of RNA binding proteins (Malhotra & Sowdhamini, 2014; Brannan *et al*, 2016; Ghosh & Sowdhamini, 2016), comprehensive overview articles (Gerstberger *et al*, 2014; Neelamraju *et al*, 2015; Sundararaman *et al*, 2016; Cook *et al*, 2011), and manual curation. For the genome-wide screening, we used the Human Genome ON-TARGETplus siRNA Library-SMARTpool (Dharmacon) that targeted 18104 genes. This library comprised three subsets: Drug Targets (G-104655-E2, Lot 11169), Druggable (G-104675-E2, Lot 11167), and Genome (G-105005-E2, Lot 11170).

### Control siRNAs

Screen progress was monitored with seven control transfections: three positive controls to monitor transfection efficiency and four negative controls to ensure our competitive growth assay was unaffected by the transfection procedure itself and cells were in good condition. The positive controls were siRNAs that targeted the EGFP coding sequence and *PLK1* and *UCB1* genes. The siRNA that targeted EGFP should cause a large decrease in the *green fraction* (*gf*) because “green” cells would lose their fluorescent reporter and become “black.” *PLK1* and *UCB1*are essential house-keeping genes. Their knockdown is lethal, and efficient transfection should greatly decrease cell number. We had no way of knowing *a priori* how it would affect *gf*. Thus, we focused on the strength of the effect and whether the effect was constant throughout the large screens.

In the genome-wide screen ofDIS3, after processing the first batch of cells, we noticed that silencing of the *PCF11* gene produces an extraordinarily low *gf* and decided to replace the *UCB1* control with *PCF11* in subsequent transfections.

For negative controls, we used the non-targeting siRNAs neg1, neg3, and neg4 and their pool (negp). Notably, the pool also contains neg2 siRNA. We decided to not use neg2 alone because it had a uniquely cytotoxic effect of considerable magnitude that was not replicated with negp.

### Plate layout

In the siRNA libraries, the siRNAs that target particular genes usually occupy rows B to O and columns 3 to 22. These are called *sample* wells. We left rows A and P empty and plated cells in all columns in rows B to O. Control siRNAs were placed in columns 2 and 23, and cells in columns 1 and 24 were left non-transfected (nt).

### Image acquisition and analysis

Imaging was performed using the ScanR system (Olympus) that was equipped with a MT20 illumination unit and 150 W mercury-xenon burner and a DAPI/FITC/Cy3/Cy5 quad Sedat filter set (Semrock) using a 20×/0.75 objective lens in the extended RNA metabolism screen and a 10×/0.35 PlnFLN lens in all of the other screens. Single images (not stacks) were taken from six fields of view in every well in the extended RNA metabolism screen and five fields of view in all of the other screens. In the genome-scale screen, the images were binned 2×2 to shorten acquisition times. ScanR Acquisition software was used for microscope control.

Primary image analysis was performed using ScanR Analysis software. Every channel was run through rolling-ball background subtraction. Segmentation was performed on the Hoechst channel using the Edge algorithm. The identified objects were gated based on a scatter plot of the circularity factor relative to the area. The objects that comprised the gate are hereinafter referred to as cells. The mean intensity of the EGFP and mCherry channels was calculated for all cells, and the two intensities were plotted on a scatter plot. Cells were classified into one of three groups: black (no reporter fluorescence), green (high EGFP fluorescence and no mCherry fluorescence), and red (high mCherry fluorescence and no EGFP fluorescence). The numbers of cells in each gate in each well were exported as tab-delimited text files.

### Metrics

In our analysis, we noted two population metrics: general viability and relative fitness. General viability is the total number of cells in a particular well compared with nt cells in the respective plate. It is calculated plate-wise as *relative viability* (Equation 1) and reflects the cytotoxicity of particular siRNAs. Relative fitness reflects the direction and strength of the genetic interaction. Relative fitness is expressed as the fraction of cells that express the WT protein of interest among all cells that expressed a transgene (i.e., the fraction of green cells among all positive [green and red] cells). We called this metric the *gf* and calculated it well-wise (Equation 2).

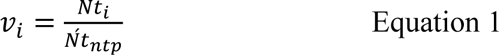

where *vi* is relative viability in well *i*, *Nt_i_* is the total number of positive cells in well *I* (green + red), and *Nt_ntp_* is the mean total number of cells in all nt wells in the respective plate *p*. Negative cells (black; see below) were omitted for the purpose of this calculation because their number was always very low (not shown).

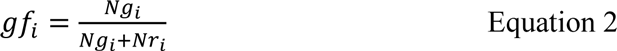

where *gf_i_* is the “green fraction” in well *i*, *Ng_i_* is the number of green cells in well *i*, and *Nr_i_* is the number of red cells in well *i*.

### Statistical analysis and hit selection

Downstream analysis was performed in the R environment (Gentleman *et al*, 2004) using home-written tools. The pipeline made use of the following packages: magrittr (Wickham, 2014), tidyr (Wickham *et al*, 2020b), dplyr (Wickham *et al*, 2020a), ggplot2 (Wickham, 2016), data.table (Dowle *et al*, 2020), and reutils (Schöfl, 2016).

Briefly, all of the result files were loaded and collated into a single data frame. *gf* was normalized by running Tukey’s median polish on each plate. The normalized values were then converted to robust *z*-scores (*z\** scores) according to Equation 3.

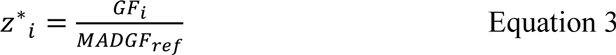

where *z*i* is the *z\** score for well *i*, *GF_i_* is the normalized *gf* in well *i*, and *MADGF_ref_* is the median absolute deviation of normalized *gf* in the reference set. *z\** scores were calculated globally and not plate-wise. The reference set in the larger screens (*i.e*., extended RNA metabolism) was the sample wells because there were many sample siRNAs, and only a few were expected to produce a reaction. In the core RNA metabolism screens, the sample siRNAs targeted a small group of hand-picked genes with probable genetic interactions, which was likely to skew the baseline; therefore, we used untreated wells as a reference for *gf*.

The *z\** score reflects the differential viability between WT and MUT cells. A positive *z\** score means that there are more WT cells than MUT cells. A negative *z\** score means the opposite. The absolute value of the *z\** score reflects the difference between WT and MUT cell viability, which translates to the magnitude of epistasis and thus to the strength of the GI between the target gene in that well and the gene of interest in the screen.

We set the *z\** score threshold for being considered a hit to a sufficiently low number to obtain a substantial amount of hits but sufficiently high to not obtain hits in nt wells and wells that were transfected with negative control siRNAs. Notably, the thresholds were set independently for each screen, ranging from 2 to 2.4.

Based on the *z\** scores, each well received a hit score: 1 if the *z\** score was higher than the threshold, −1 if the *z\** score was lower than the negative threshold, and 0 if the *z\** score fell between a negative and positive threshold. Hit scores were then summed across replicates to yield summarized hit scores. Wells with an absolute summarized hit score of 2 or more were considered hits in their respective screens. However, wells with relative viability less than 30% were disregarded when summarizing hit scores.

### RNA-Seq

#### Cell culture

Stably transfected derivatives of HEK293 FLP-In T-Rex cells that expressed WT DIS3 or the catatonically inactive version (G766R) of DIS3 were cultured for transfection in six-well format in DMEM (Gibco) supplemented with 10% FBS (Gibco) without antibiotics at 37°C in a 5% CO_2_ humidified atmosphere.

The expression of transgenes that coded for sh-miRNA-EGFP cassettes to silence endogenous DIS3 and sh-miRNA-insensitive FLAG-tagged exogenous DIS3 protein was induced with tetracycline at a final concentration of 25 ng/ml 24 h before siRNA transfection. For siRNA-mediated SF3A1 depletion, tetracycline-induced cells were subjected to Silencer Select siRNA s20118 transfection (Ambion, catalog no. 4392421). Scrambled Silencer Select Negative Control siRNA (Ambion, catalog no. 4390844) was used as a negative control. Transfections were performed using Lipofectamine RNAiMAX (Invitrogen, catalog no. 13778-150) and 20 nM siRNA according to the manufacturer’s recommendations and cultured for ∼72 h before harvesting. The RNA-Seq experiments were performed in duplicate.

#### Library preparation and sequencing

Total RNA was isolated with TRI Reagent (Sigma-Aldrich, catalog no. 93289) according to the manufacturer’s instructions. Following DNase treatment, strand-specific RNA libraries were prepared using a KAPA Stranded RNA-Seq Library Preparation Kit (KAPA Biosystems, catalog no. KK8401) according to the manufacturer’s protocol. The libraries were sequenced using an Illumina NextSeq500 sequencing platform in the 150-nt paired-end mode.

#### Quality control and mapping of next-generation sequencing data

The Illumina sequencing reads were quality filtered using Cutadapt 1.18 (Martin, 2011) to remove Illumina adapter sequences, trim low-quality fragments (minimum Q score = 20), and remove reads that were shorter than 30 nt after trimming. For JUM alternative splicing analysis, reads were mapped to the human genome (hg38) using STAR 2.6.0c (Dobin & Gingeras, 2015) with the two-pass mode for better junction discovery. Before mapping, the genome was indexed with Gencode v28 basic annotation. For downstream analysis, only uniquely mapped reads were used.

### Differential alternative splicing analysis

Alternative splicing (AS) events were analyzed using JUM (Wang & Rio, 2018). For AS structure quantification, JUM uses RNA-Seq reads that are mapped to splice junctions. Only splice junctions that received more than five reads in each of the replicates of the siRNA-treated and control samples were considered valid junctions for downstream analysis. After profiling AS structures in each condition, JUM performs an analysis of differential AS structure usage between examined conditions. As a metric for quantifying differential splicing events between conditions, JUM calculates the ΔPSI. PSI represents the percentage of transcripts, including a particular exon or splice site, and is calculated from the read that is mapped to splice junctions. For the quantification of changes in AS, JUM compares the usage of each profiled AS by modeling the total number of reads that are mapped to AS sub-junctions with a negative binomial distribution and evaluating the overall change in the basal expression of AS between conditions. Only AS events that showed more than 10% of a difference (|ΔPSI|> 0.1) were considered differentially spliced. An Benjamini–Hochberg-adjusted *p* value of 0.05 was used as the statistical threshold for significance.

### Gene Ontology enrichment analysis

The GO analyses were performed in the R/Bioconductor environment (Gentleman *et al*, 2004; Huber *et al*, 2015) using the clusterProfiler package (Yu *et al*, 2012) and data from the Database for Annotation, Visualization and Integrated Discovery (DAVID; https://david.ncifcrf.gov/home.jsp; Huang *et al*, 2009) with the RDAVIDWebService package (Fresno & Fernández, 2013). As a background for enrichment analysis, we used lists of genes that were tested in each of the analyzed libraries. The minimum number of genes in enriched processes was set to 2. The analyses were performed based on annotations of Biological Processes, Molecular Functions, and Cellular Compartments. The results are presented in the Datasets EV3, EV5, EV7. The visualization of enrichment analysis was performed by plotting bar plots that represented the number of genes that were enriched in the top 10 categories, sorted by a Fischer’s exact test *p* value using the ggplot2 package.

### Gene Ontology enrichment network analysis

The GO enrichment networks were created in the R/Bioconductor environment using the igrpah package, ggraph package, and home-written tools. To evaluate relationships between distinct biological processes from the GO analyses, we created networks that represented connections between these processes with shared genes that were assigned to them. The list of enriched functional categories and genes were extracted for the enrichment results and converted to graph format (igraph::graph.data.frame). In-network construction connections between genes and biological processes, represented as edges, were created based on the assignment of genes to extracted biological processes according to the DAVID GO library, with genes and biological processes as nodes. The networks were visualized with ggraph. The networks are presented in the Fig. 2, 3, EV3. Figures EV4 and EV5 were constructed so that edges represent the number of genes that are commonly assigned to connected functional categories that constitute nodes in those graphs.

### Genetic Interaction Network Generation

The resulting significant hits from the core RNA metabolism high-throughput siRNA screen were visualized in the network layout using Cytoscape (Shannon *et al*, 2003). For visualization, networks were defined as a list of nodes that represented query ribonucleases and genes that were defined as significant hits in the screen. The connection between nodes was based on the resulting genetic interaction that was identified in the screen and classified into two possible categories: synthetic lethal (blue edges) and suppressor (yellow edges). Additionally, all hits were analyzed in the STRING protein-protein (P-P) association database (Szklarczyk *et al*, 2019) to investigate overlap between known P-P interactions and identified genetic interactions. The extracted P-P interactions between query genes and hits are represented in the network with gray edges. To visualize interactions with known protein complexes, genes that interacted with screen hits and are known to form protein complexes were grouped together.

## DATA AVAILABILITY

The datasets and computer code produced in this study are available at:

- RNA-Seq data: Gene Expression Omnibus GSE155123 (https://www.ncbi.nlm.nih.gov/geo/query/acc.cgi?acc=GSE155123)
- Package for screen data analysis (development version): GitHub (https://github.com/olobiolo/siscreenr)

## ACKNOWLEDGEMENTS

We acknowledge former and current members of AD lab for stimulating discussions and all support; Katarzyna Prokop and Dorota Adamska for technical support in NGS experiments; Michal Malecki for helpful discussions and suggestions on screening analyses. Financial support was provided by the TEAM program of the Foundation for Polish Science, financed by the European Union via the European Regional Development Fund (agreement no. TEAM/2016-1/3 to AD) and ERA Chair European Union’s Horizon2020 program (agreement no.810425).

## AUTHOR CONTIBUTIONS

AD and RJS developed and directed the studies. AH-O set up the procedure of computational and bioinformatic analysis of normalized siRNA screen results, performed GSEA and network analyses and carried out RNA-Seq data analyses, wrote the manuscript, and prepared all figures and tables with feedback from AD. AC performed high-throughput microscopy, image analysis, designed and programmed the siRNA screening analysis, and participated in writing the manuscript. KA, KK, PO contributed to the technical component of screening implementation: established stable cell lines, performed cell culturing and transfections. RJS designed the siRNA screening procedure and subgenomic siRNA libraries, performed experiment with depletion of splicing component, participated in writing the manuscript. AD provided funding and revised the manuscript.

## SUPPLEMENTARY FIGURE LEGENDS

**Figure EV1.**
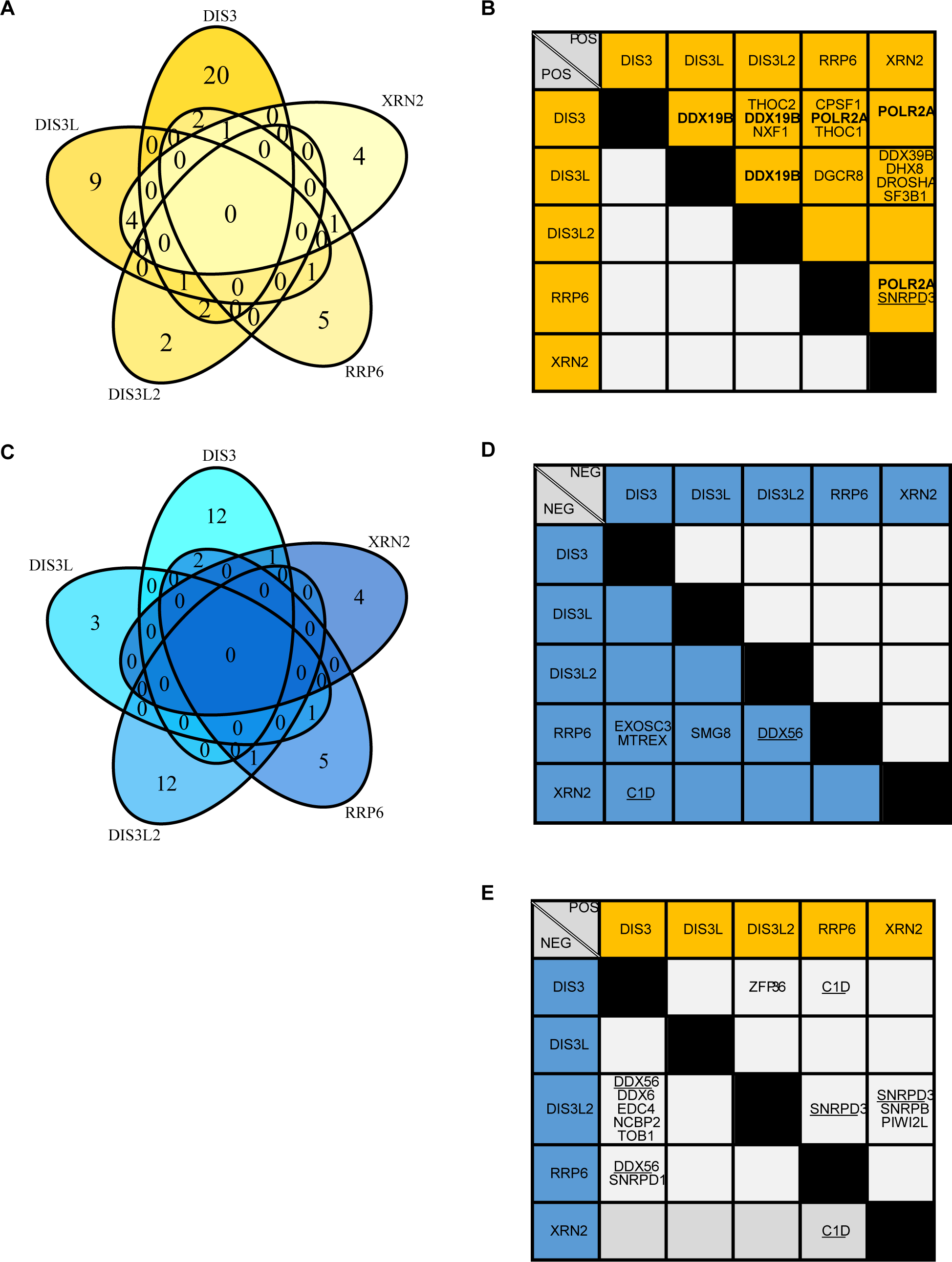
Summary of overlap of synthetic lethal and suppressor interactions with main mammalian RNases identified in the siRNA screen with a custom core RNA metabolism siRNA library. A, C. Venn diagrams representing shared and unique genetic interactions for DIS3, DIS3L, DIS3L2, EXOSC10 and XRN2. The overlap was analyzed separately for synthetically lethal (A) and suppressor (C) interactions. B, D, E. Tables summarizing common interactions for DIS3, DIS3L, DIS3L2, EXOSC10 and XRN2 identified in custom core RNA metabolism siRNA screen. The tables were divided to summarize common synthetic lethal interactions (B), suppressors (D), and interactions that were opposite for both ribonucleases (E). Genes showing the same genetic interactions with three ribonucleases are in bold, and genes that interact with three nucleases but in a complex way are underlined.

**Figure EV2.**
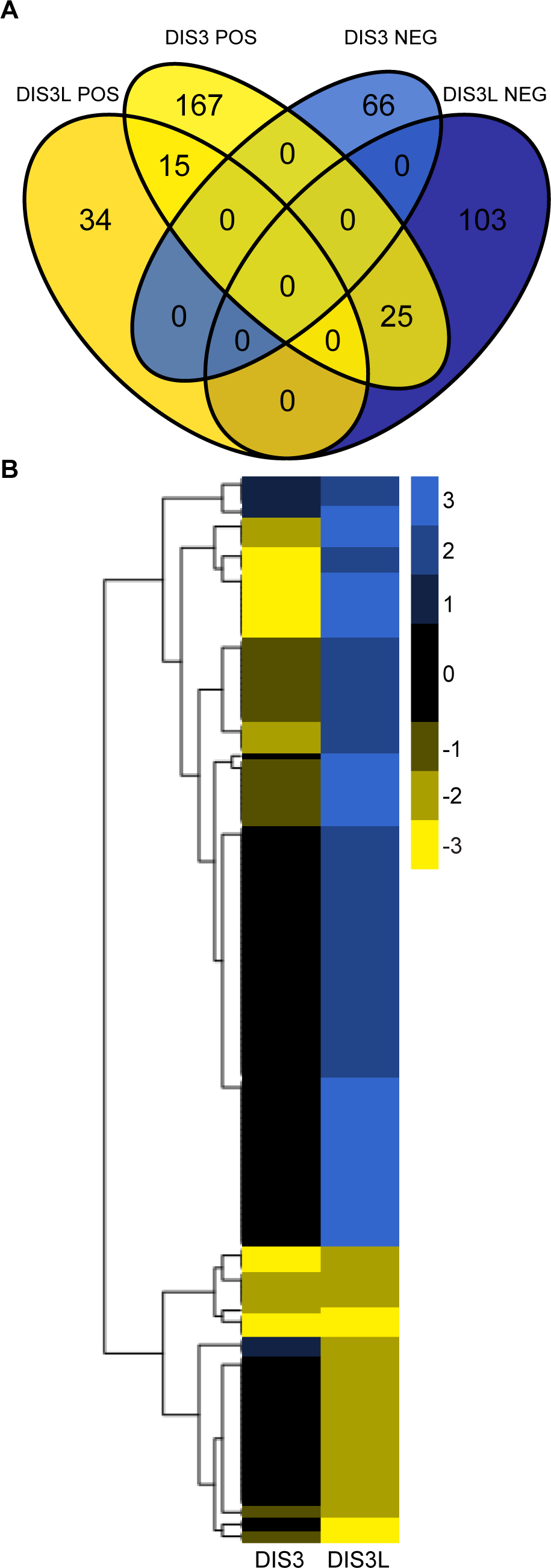
Summary of overlap of synthetic lethal and suppressor interactions with DIS3 and DIS3L identified in the siRNA screen with a custom RNA metabolism extended siRNA library. A. Venn diagrams show shared and unique genetic interactions for DIS3 and DIS3L. B. Heatmap representation of summarized z* scores of genetic interactions between DIS3 and DIS3L and RNA metabolism genes. The plots represent scores for 410 of 3904 tested genes that were identified as a significant hit for DIS3 or DIS3L. Yellow represents positive hits (suppressors). Blue represents negative hits (synthetic lethal). Black represents genes that have no interactions.

**Figure EV3.**
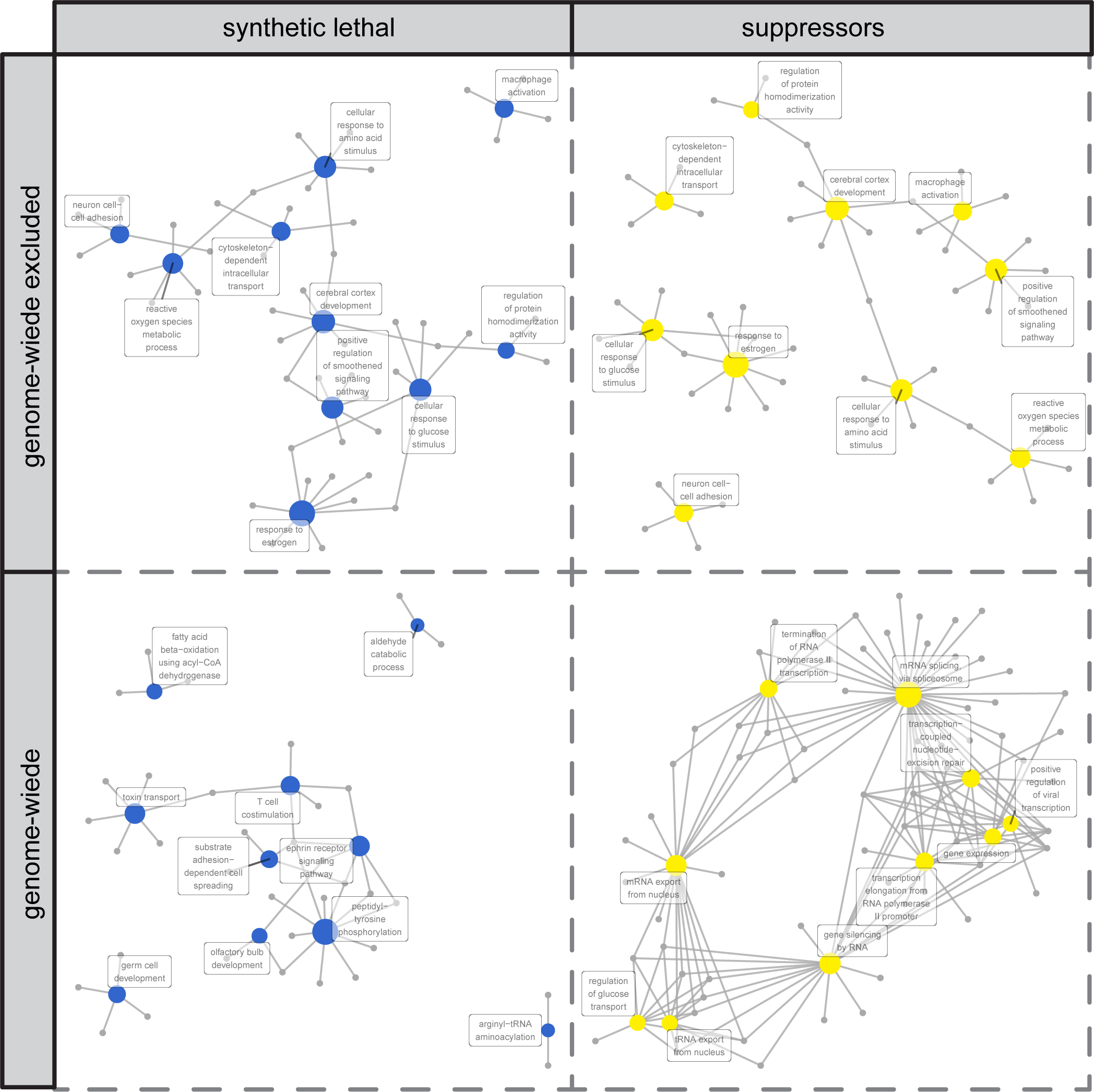
DIS3 genetic interactions are mainly involved in RNA metabolism processes. Networks illustrate interactions between biological processes that share genes that were identified in the genome-wide siRNA screen of DIS3 and in the subset of hits from the genome-wide screen that excluded hits identified in the extended RNA metabolism screen. The colored nodes (blue and yellow) represent biological processes that were interconnected and shared the same genes that were identified as synthetic lethal interactions or suppressors, respectively. The networks were constructed in R/Bioconductor based on the associations of hit genes with biological processes according to the DAVID direct GO database.

**Figure EV4.**
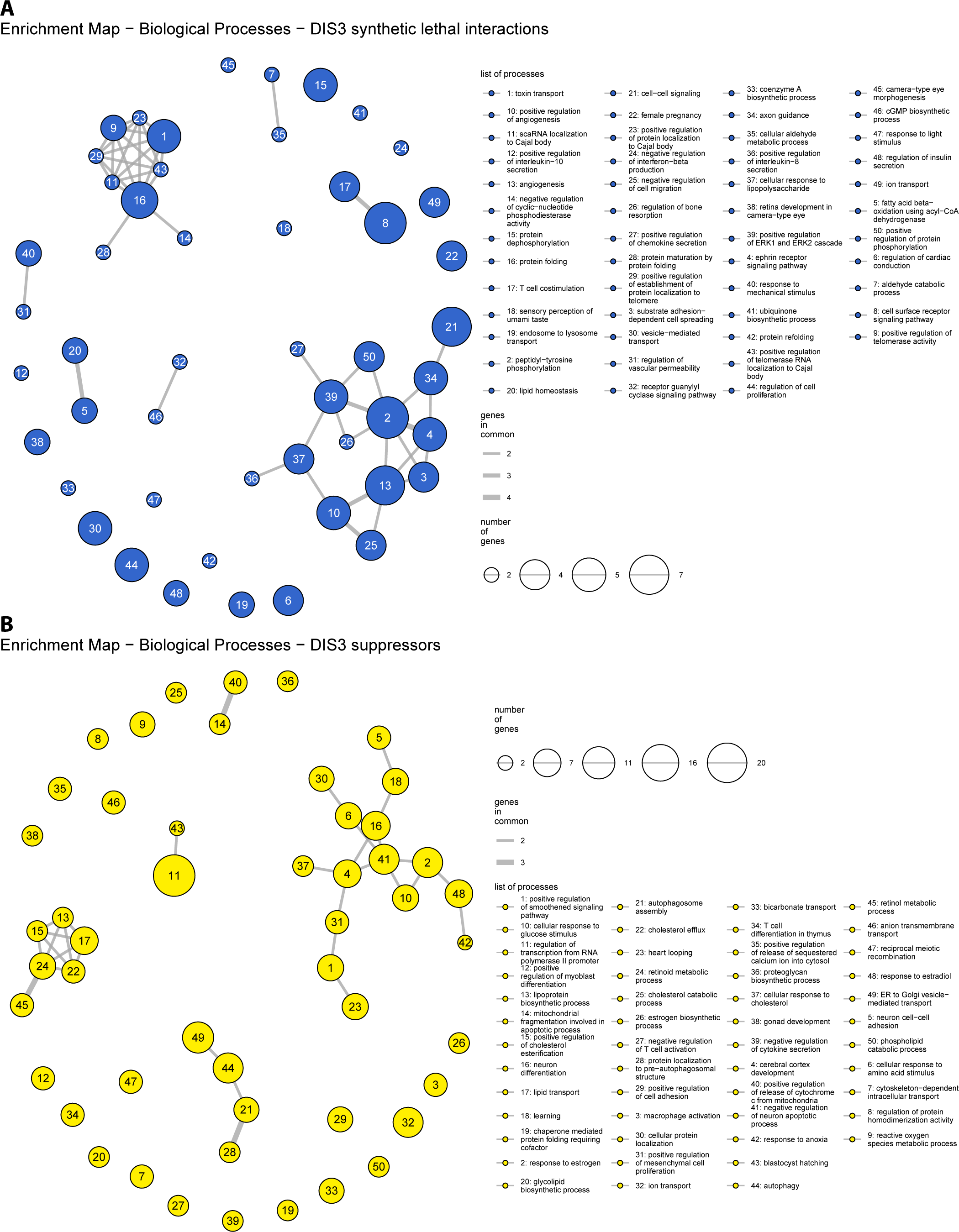
Enrichment map of Biological Processes associated with genes identified as genetic interactors of DIS3. A-B. The maps display clusters of biological processes enriched by genes that were identified as (A) synthetic lethal interactions and (B) suppressors of DIS3. The enrichment was performed on the subset of hits that were identified in genome-wide screen, excluding hits identified in the extended RNA metabolism screen. The networks were constructed in R/Bioconductor based on associations of hit genes with biological processes according to the DAVID direct GO database.

**Figure EV5.**
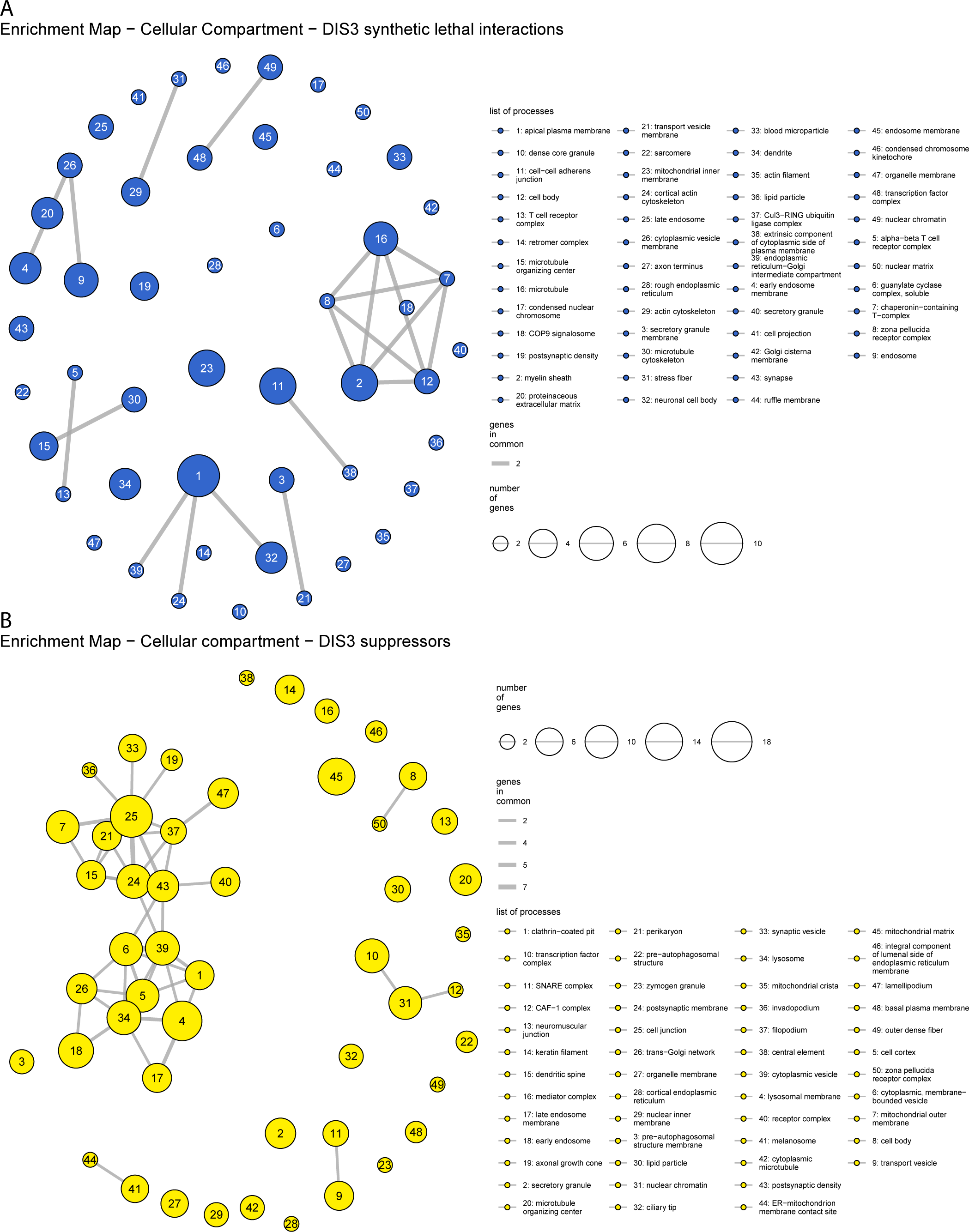
Enrichment map of Cellular Components associated with genes identified as genetic interactors of DIS3. A-B. The maps display clusters of cellular components that were enriched by genes identified as (A) synthetic lethal interactions and (B) suppressors of DIS3. The enrichment was performed on the subset of hits that were identified in the genome-wide screen, excluding hits identified in the extended RNA metabolism screen. The networks were constructed in R/Bioconductor based on the associations of hit genes with biological processes according to the DAVID direct GO database.

